# *In vivo* deuteration reveals pronounced variation in myelin lipid turnover rates and reduced myelin renewal with ageing

**DOI:** 10.64898/2026.02.17.706445

**Authors:** Jun Yup Lee, Yushan Cai, Mika Westerhausen, Jesse A. Michael, Jonathan D. Teo, Huitong Song, Georgia Watt, Shane R. Ellis, Anthony S. Don

## Abstract

Myelin turnover is essential for its structural and functional integrity, yet how this particularly lipid-rich membrane is renewed and why it deteriorates with ageing remain unresolved. Combining deuterium oxide administration in mice with high resolution lipidomics, we establish that brain lipid turnover rates are highly heterogeneous, differ by brain region, and depend primarily on lipid class. Half-lives of common glycerophospholipids in purified myelin were under 2 months whereas many sphingolipids exhibited half-lives exceeding 8 months, dependent on acyl chain length and saturation. Myelin sphingolipid and cholesterol replacement rates in the corpus callosum decreased markedly between 3 and 12 months of age, while disrupting lipid trafficking through ApoE ablation preferentially impaired cholesterol turnover and incorporation into myelin. Our results establish that myelin renewal occurs through continual replacement of individual lipid constituents in a manner that depends on lipid class, hydrophobicity, and ApoE-dependent trafficking, and that this process slows significantly with ageing.

## Introduction

Brain white matter volume decreases significantly from middle age, a process that accelerates with ageing, whereas grey matter volume declines at a more modest rate throughout the adult lifespan^1,2^. This loss of white matter volume is very likely a consequence of myelin loss and is associated with reduced executive function, memory, and walking speed with ageing^3–5^. Oligodendrocytes and myelin are essential not only for rapid saltatory conduction along neuronal axons, but also for provision of trophic and metabolic support to neurons^6^. Myelin degeneration promotes amyloid-beta formation^7^, and age-dependent myelin deterioration is likely to be a significant pathogenic driver for Alzheimer’s disease and frontotemporal dementia^8–10^.

^14^C tracing in post-mortem human corpus callosum samples indicates that myelin undergoes constitutive turnover while the oligodendrocyte population remains stable throughout adult life^7^. In fact, continuous myelin synthesis and degradation is essential for its integrity, as gene knockouts that prevent the synthesis or autophagic degradation of myelin in adult mice cause degeneration of myelin and neuronal axons^11–13^. Oligodendrocytes are known to catabolise myelin proteins through autophagy under normal physiological conditions^12,13^. Microglia and astrocytes also phagocytose and catabolise myelin lipids and proteins during development, ageing, or under pathological conditions in which there is myelin degeneration^14–17^. A key question in the field is whether constitutive myelin turnover occurs through the internalisation, breakdown and replacement of whole myelin membrane segments, or continuous replacement of individual lipid and protein constituents^18^.

Myelin is comprised 70-80% of lipid (dry weight), making it a particularly lipid-rich membrane^19,20^. Accordingly, the brain is one of the most lipid-rich organs and many neurological diseases are associated with defects in brain lipid synthesis, catabolism, or transport^21–23^. Myelin also bears a distinct lipid composition, with a high proportion of cholesterol, hexosylceramide (HexCer) and its sulfated derivative sulfatide (SHexCer), and phosphatidylethanolamine (PE) plasmalogens^19,20^. The distinctive lipid composition of myelin is essential for its stability. For example, in the CNS, over 99% of HexCer is galactosylceramide (GalCer)^24,25^, which accounts for 20-25% of myelin lipid^19,20^. Defects in GalCer and SHexCer synthesis or acyl chain length destabilise myelin and cause neurodegeneration^24,26^. Similarly, defects in lysosomal catabolism of GalCer and SHexCer cause the neurodegenerative diseases Krabbe’s disease and metachromatic leukodystrophy^23^.

Developments in mass spectrometry over the past two decades have provided the means to rapidly quantify hundreds of lipids, however conventional lipidomics only affords a snapshot of the lipidome at a specific point a time. Methods that quantify rates of brain lipid synthesis and turnover will provide powerful insights into brain physiology and the genetic basis for many neurological diseases. Global protein synthesis and turnover have been quantified by incorporation of ^13^C-labelled lysine or ^15^N-labelled algae into mouse feed, leading to the observation that myelin proteins are very long-lived^27–29^. ^13^C-labelled fatty acids, carbohydrates, or acetate can be used to track lipid metabolism but introduce bias due to their differential incorporation into lipids based on biosynthetic pathways and metabolic status^30^. In contrast, administration of deuterium oxide (^2^H_2_O) can result in unbiased incorporation of the stable hydrogen isotope deuterium (^2^H) into lipids^31–33^.

This study combined ^2^H_2_O administration in the drinking water with high resolution mass spectrometry to quantify turnover rates for a wide range of glycerophospholipids, sphingolipids, and cholesterol in dissected brain tissue and purified myelin. This revealed pronounced variation in membrane lipid turnover rates, whereby common cellular glycerophospholipids were almost fully replaced within 8 weeks, whereas myelin-enriched sphingolipids turned over very slowly, with half-lives exceeding those of long-lived myelin proteins. Turnover of these long-lived sphingolipids and cholesterol was significantly reduced with ageing, and this was exacerbated by absence of the major CNS lipid carrier apolipoprotein E (ApoE), supporting a model in which individual lipid constituents of myelin are replaced through distinct mechanisms, rather than as whole myelin sheaths.

## Results

### Quantification of brain lipid turnover *in vivo*

To label brain lipids with ^2^H, mice were administered ^2^H_2_O in the drinking water from embryonic day 14 to post-natal day 60 and deuteration of each lipid was followed at 0, 2, 6, and 10 months after removal of ^2^H_2_O using high resolution mass spectrometry (Fig. 1A,B). Isotopologue peaks (i.e. the same lipid bearing different numbers of ^2^H atoms) were quantified using a semi-automated workflow, with computational correction for naturally occurring isotopes. Example spectra for the myelin lipid HexCer(d18:1/24:0) are shown in Fig 1C, and the isotopologue distribution patterns for six example lipids after de-isotoping are shown in Fig. 1D. Quantification of deuterated isotopologues was restricted to lipids for which there were no co-eluting, interfering ions in samples from unlabelled (i.e. no ^2^H_2_O) control mice following isotope correction. At the end of the developmental labelling period, >99% of common phospholipid and sphingolipid molecules, and >90% of cholesterol, carried one or more ^2^H atoms. The extent of deuteration decreased over time following removal of ^2^H_2_O from the drinking water, reflecting the turnover or replacement of deuterated lipids with newly synthesised, unlabelled lipids. The isotopologue distribution at each timepoint was quantified as the weighted mean deuteration state, which was used to determine the percentage loss of deuteration for each lipid relative to the 2-month maximal labelling timepoint (Fig. 1E). This followed a one-phase exponential decay for most lipids, permitting the calculation of lipid deuteration half-lives.

**Figure 1.**
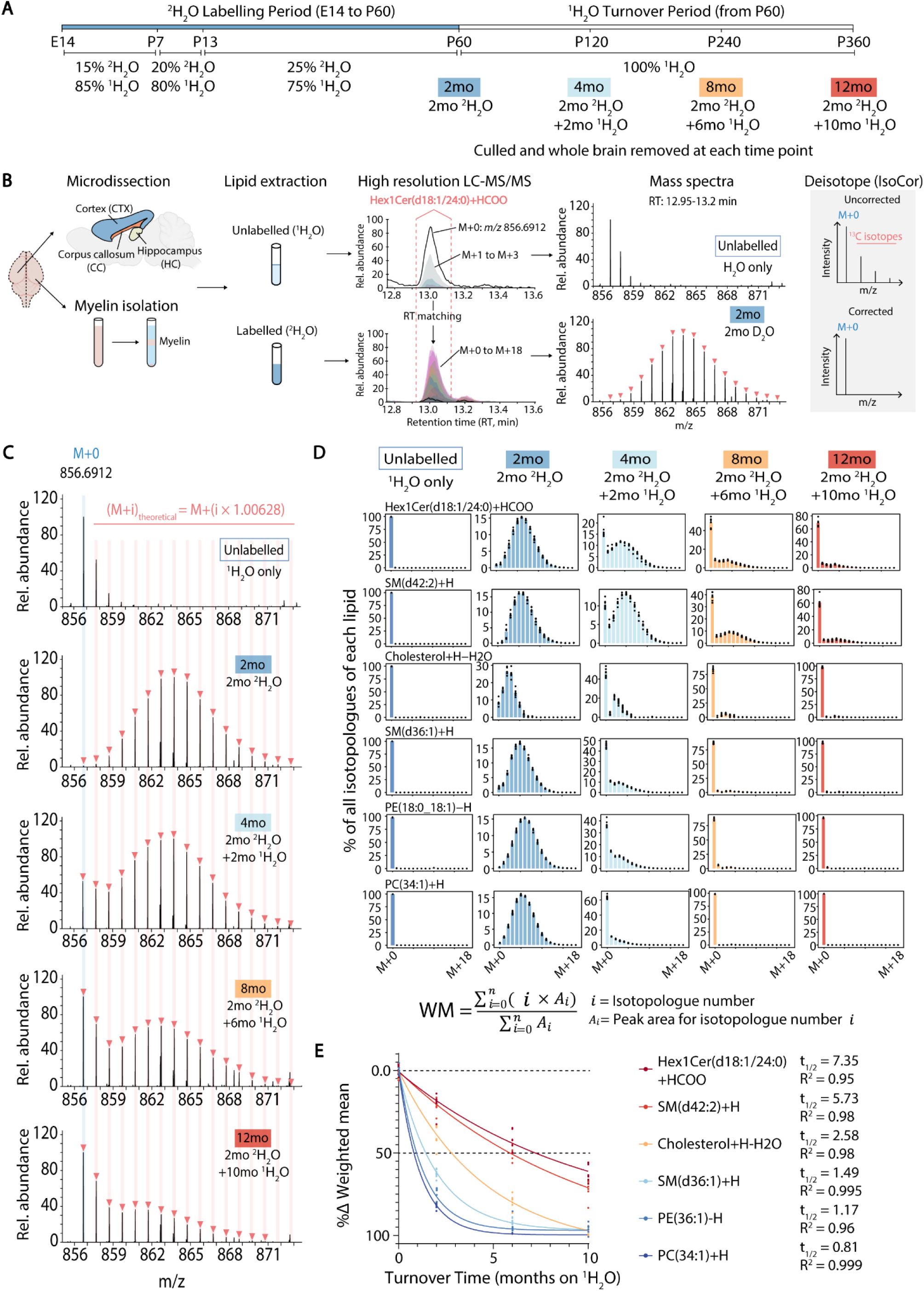
Developmental ^2^H_2_O administration results in extensive deuteration of brain lipids and allows their turnover to be followed over time. (A) Experimental timeline for ^2^H_2_O administration in the drinking water, followed by turnover time with only ^1^H_2_O. 2mo: 2-month. (B) Workflow for detection and quantification of deuterated lipid isotopologues. Lipids were identified using matched precursor and product ions in samples from mice given only ^1^H_2_O, after which deuterated isotopologues of each lipid in mice given ^2^H_2_O were identified on the basis of accurate precursor *m/z* and HPLC retention time (RT). Peak areas were then corrected for the expected abundance of naturally occurring isotopes, particularly ^13^C. (C) Raw mass spectra of myelin-enriched lipid Hex1Cer(d18:1/24:0) from mice culled after 2 months of ^2^H_2_O administration (blue), 2 months ^2^H_2_O followed by 2 months ^1^H_2_O (cyan), 2 months ^2^H_2_O followed by 6 months ^1^H_2_O (orange), and 2 months ^2^H_2_O followed by 10 months ^1^H_2_O (red). Spectra corresponding to a 0.1 min LC-MS/MS peak are shown. (D) Isotopologue distribution of Hex1Cer(d18:1/24:0), SM(d42:2), cholesterol, SM(d36:1), PE(18:0_18:1), and PC(34:1) following computational correction for naturally occurring isotopes. Distributions are shown at 0 (n=3 females, n=3 males), 2 (n=5 females, n=3 males), 6 (n=5 females, n=2 males), or 10 (n=4 females, n=3 males) months of turnover time (on ^1^H_2_O). (D) Change in the weighted mean deuteration state (WM) as a function of turnover time for lipids shown in (C), i = isotopologue number, A_i_ = peak area for that isotopologue. WM as a function of turnover time followed an exponential decay pattern. Deuteration half-lives and R^2^ values for each lipid are indicated below.

### Myelin-enriched glycosphingolipids exhibit much longer half-lives than glycerophospholipids

Deuterated isotopologues of 209 lipid ions were quantified, from which the turnover rates of 119 brain lipids were calculated (Fig. 2A; Supplementary Data File 1). The longest-lived lipids in the corpus callosum – a white matter tract that connects the hemispheres – were sphingolipids [HexCer, SHexCer, and sphingomyelin (SM)], with deuteration half-lives exceeding 4 months. Cholesterol and PEs displayed intermediate half-lives in the range 2-4 months, whereas the glycerophospholipids phosphatidylcholine (PC), phosphatidylinositol (PI), phosphatidylglycerol (PG), and phosphatidylserine (PS), and the lysophospholipids (LPC, LPE, LPS) generally had half-lives of less than 2 months. Interestingly, deuteration half-lives for HexCer and SHexCer were dependent on the length and saturation of their N-acyl chains, with saturated and very long chain species turning over more slowly than unsaturated and long chain species (Fig. 2A). In contrast, the presence of a 2-hydroxyl group in the N-acyl chain (e.g. d18:1/24:1(OH)) did not have any appreciable effect on half-life.

**Figure 2.**
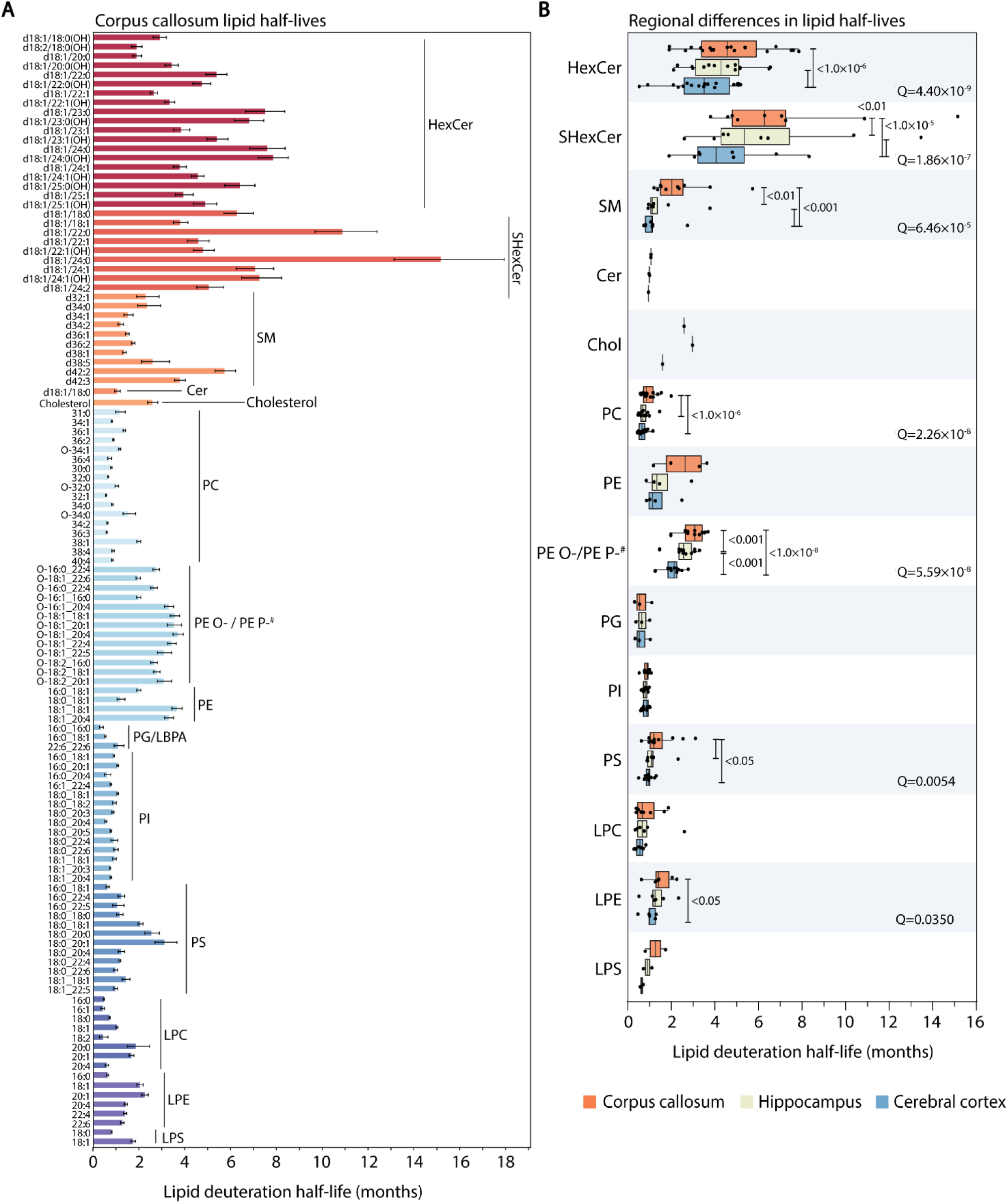
Lipid half-lives vary by functional class, acyl chain length, and saturation state. (A) Half-lives of individual lipid species in the corpus callosum of C57BL/6 mice (2mo: n=3 females, n=3 males; 4mo: n=5 females, n=3 males; 8mo: n=5 females, n=2 males; 12mo: n=4 females, n=3 males). Errors bars represent 95% confidence interval. (B) Repeated measures ANOVA was used to compare turnover rates of each lipid class between the corpus callosum (orange), hippocampus (green), and cerebral cortex (blue). Each datapoint represents a distinct lipid. Q values are shown for lipid classes that differed significantly between regions after adjusting for multiple comparisons, and adjusted P values from Tukey’s post-hoc test are displayed for relevant comparisons. HexCer: hexosylceramide, SHexCer: sulfatide, SM: sphingomyelin, Cer: ceramide, Chol: cholesterol, PC: phosphatidylcholine, PE: phosphatidylethanolamine, PE O-/PE P-^#^: ether-linked phosphatidylethanolamine, PG/LBPA: phosphatidylglycerol, PI: phosphatidylinositol, PS: phosphatidylserine, LPC: lysophosphatidylcholine, LPE: lysophosphatidylethanolamine, LPS: lysophosphatidylserine. ^#^Isomer pairs of plasmanyl (PE O-) and plasmenyl (PE P-) PE species could not be distinguished by accurate mass, class-specific fragments, or retention time. Thus, listed PE O-species may be PE P-species with one less double bond, and vice-versa.

Lipid turnover rates differed between brain regions, with significantly faster turnover of HexCer, SHexCer, SM, PC, ether-linked PE (PE O- and PE P-)^#^, LPE, and PS lipid classes in the cortex compared to the corpus callosum (Fig. 2B). Turnover rates in the hippocampus were between those of the cortex and corpus callosum.

While HexCer and SHexCer are highly enriched in myelin^19,20^, the fast-turnover lipids PC and PI are ubiquitous to all cells, implying that myelin lipids turn over more slowly than other membrane lipids. Deuterated isotopologues of HexCer(d42:1(OH)) and HexCer(d42:2) were readily detected with mass spectrometry imaging (MSI) and localised to white matter regions (Fig. 3A,B; Extended Data Fig. 1). These deuterated isotopologues were not observed in ^1^H_2_O control mice. In agreement with our LC-MS/MS data, deuterated isotopologues of these lipids were still present in white matter-rich brain regions six months after ^2^H_2_O withdrawal (Fig. 3A,C). Ether-linked PE species, particularly plasmalogens (PE P-), are also enriched in white matter^34^ and myelin^19^, as shown in images for PE(P-34:1)/PE(O-34:2)^#^ (Fig. 3D,E). However, the deuterated isotopologues of PE(P-34:1)/PE(O-34:2) were almost fully replaced by the non-deuterated form six months after ^2^H_2_O withdrawal (Fig. 3F), demonstrating faster turnover than HexCer. In agreement with our LC-MS/MS data, deuterated isotopologues of the grey matter-localised lipids PI(38:4) (Fig. 3G-I), PE(40:6), and PS(40:6) (Extended Data Fig. 2) were fully replaced by the non-deuterated (M+0) form six months after ^2^H_2_O withdrawal.

**Figure 3.**
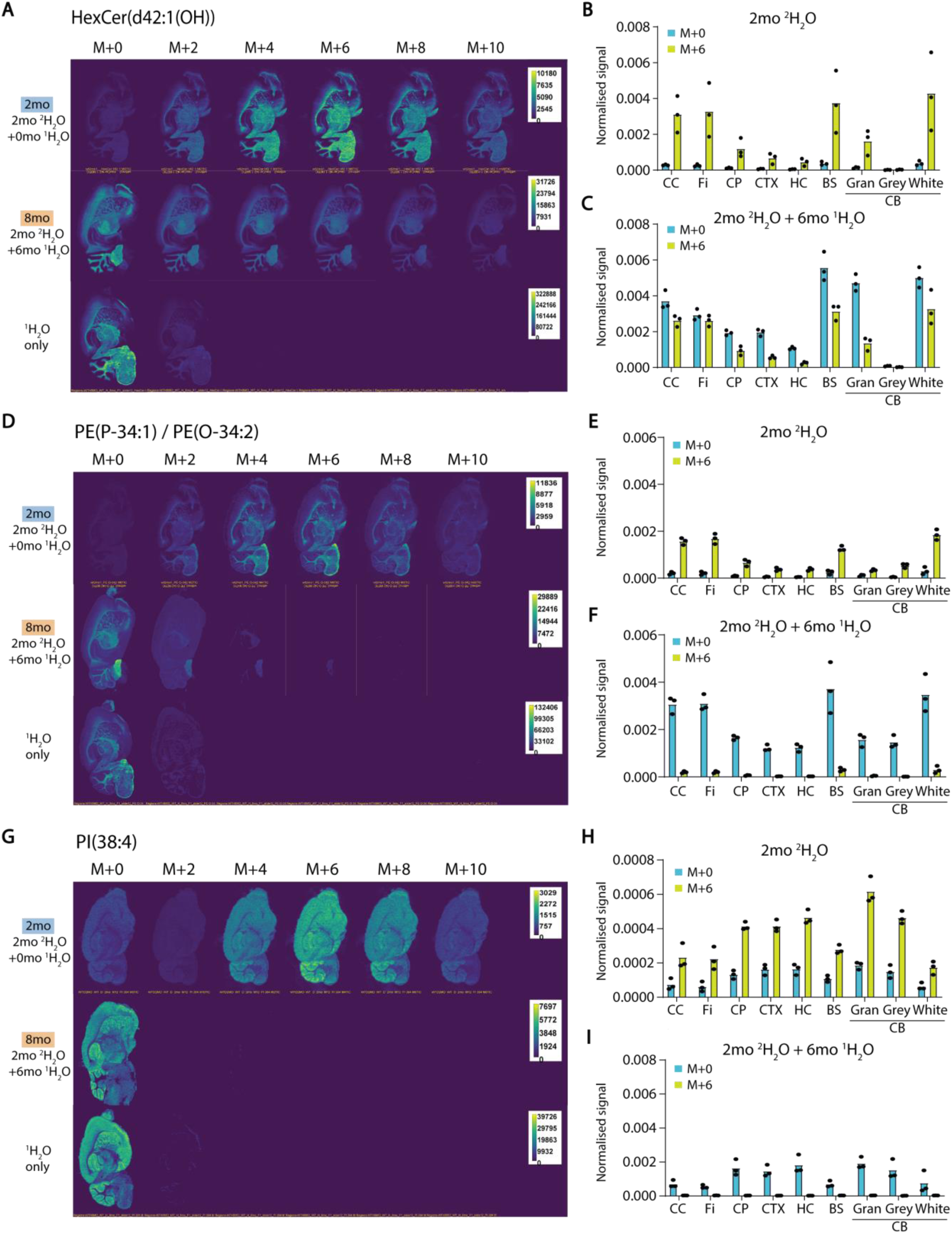
Turnover rates for white matter lipids differ according to lipid class and brain region. (A) Representative mass spectrometry images for the [HexCer(d42:1(OH))+H]^+^ monoisotope (M+0) and its deuterated isotopologues (M+2 to M+10) in mice given ^1^H_2_O only (n=3), ^2^H_2_O for 2 months (2mo, n=3), or ^2^H_2_O for 2 months followed by ^1^H_2_O for 6 months (2mo ^2^H_2_O + 6mo ^1^H_2_O, n=3). (B, C) Mean M+0 and M+6 isotopologue intensities, normalised to the total ion chromatogram, in defined brain regions of mice given (B) 2mo ^2^H_2_O, or (C) 2mo ^2^H_2_O + 6mo ^1^H_2_O. (D-F) Representative images and normalised ion intensities for [PE(P-34:1)/PE(O-34:2)+H]^+^. (G-I) Representative images and normalised ion intensities for [PI(38:4)+H]^+^. Normalised signals were averaged over defined brain regions: CC, Corpus callosum; Fi, fimbria; CP, caudate putamen; CTX, cerebral cortex; HC, hippocampus; BS, brain stem; CB, cerebellum; Gran, cerebellar granular layer; Grey, cerebellar grey matter; White, cerebellar white matter.

We also noted that the spatial distribution of HexCer(d42:1(OH)) and HexCer(d42:2) that were newly-synthesised in the 6 months after cessation of ^2^H_2_O administration differed considerably from the distribution of these same lipids at two months of age, with a much larger signal for newly-synthesised, non-deuterated lipid (M+0) relative to deuterated lipid (M+6) in the cortex and hippocampus (Fig. 3A-C and Extended Data Fig. 1). This implies relatively greater synthesis of new myelin in these regions compared to the corpus callosum, fimbria, brain stem, and cerebellar white matter between 2 and 8 months of age. In agreement with this, mean levels of myelin basic protein (Mbp) and myelin oligodendrocyte glycoprotein (Mog) increased 3-fold and 2-fold, respectively, in the cortex of untreated C57BL/6J mice between 2 and 8 months of age (Mbp p=0.0039, Mog p<0.0001), and 2-fold in the hippocampus (Mbp p=0.0004, Mog p=0.00005), but only 1.2- and 1.6-fold in the corpus callosum (Mbp p>0.05, Mog p=0.0037) and were unchanged in the cerebellum (both p>0.05) (Fig. 4A,B).

**Figure 4.**
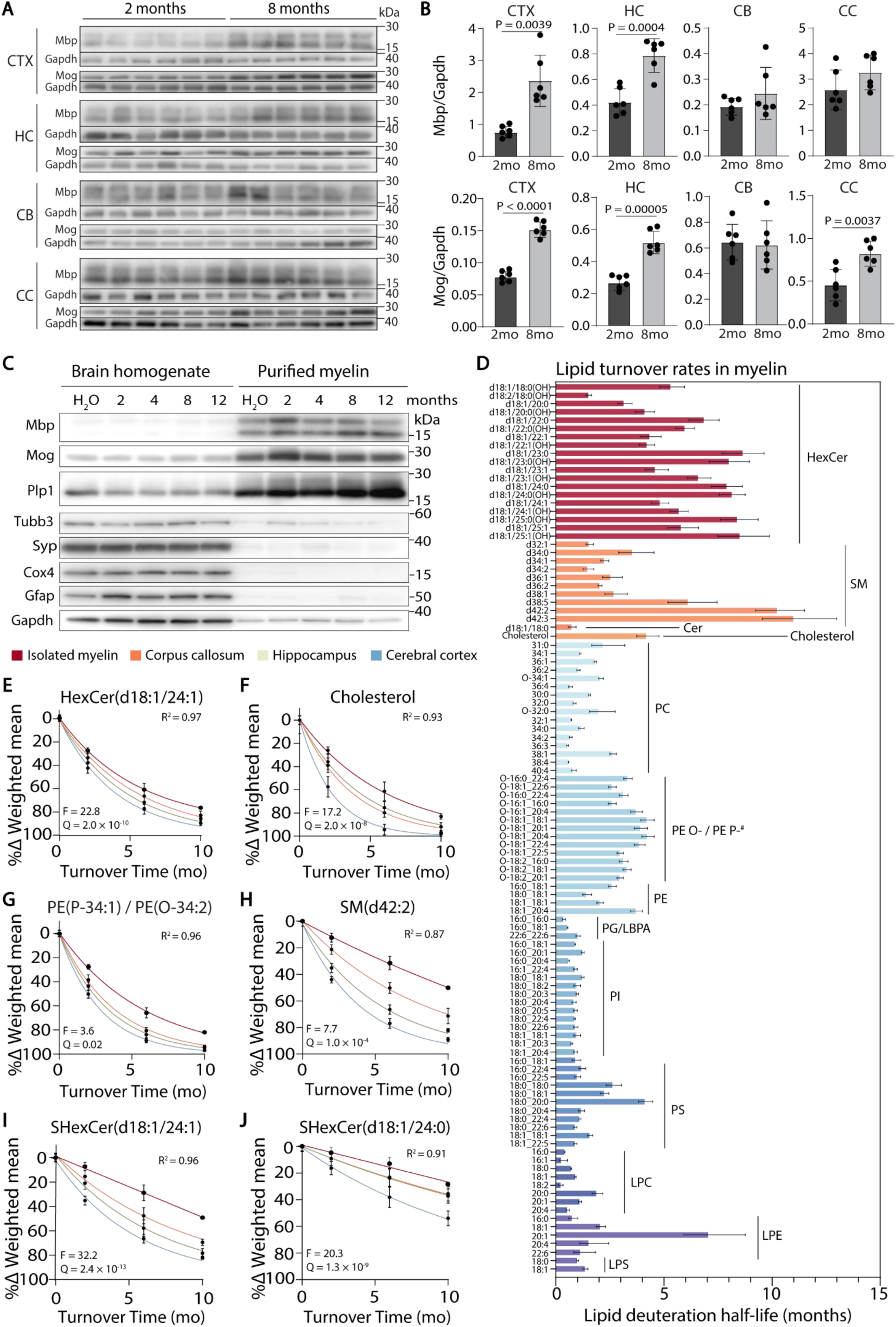
Heterogeneity in lipid turnover is preserved in isolated myelin. (A) Western blots and (B) densitometric quantification of myelin basic protein (Mbp) and myelin oligodendrocyte glycoprotein (Mog), normalised to glyceraldehyde-3-phosphate dehydrogenase (Gapdh), in 2- and 8-month old mice (3 male and 3 female per group). (C) Western blots for protein markers of myelin (Mbp, Mog, Plp1), neuronal cytoskeleton (βIII-tubulin (Tubb3)), synapses (synaptophysin (Syp)), mitochondrial inner membrane (Cox4), astrocyte cytoskeleton (Gfap), and cytosol (Gapdh), demonstrating myelin enrichment in representative mice from the 2-, 4-, 8-, and 12-month-old ^2^H_2_O sample groups, and ^1^H_2_O control group. (D) Mean lipid half-lives in purified myelin samples (3 mice per time point), with error bars indicating 95% confidence intervals. (E-J) Decay curves showing % decrease in the weighted mean deuteration state of selected lipid species in dissected corpus callosum (orange), hippocampus (green), cortex (blue), and purified myelin (red). Error bars indicate 95% confidence intervals.

### Different myelin lipids are turned over at different rates

The much slower turnover of HexCer and SHexCer compared to myelin-enriched PE species, and differential rates of turnover for different HexCer and SHexCer species, suggested that different lipids are replaced at different rates in myelin membranes. Autophagic turnover of whole myelin membranes by oligodendrocytes or endolysosomal degradation by other CNS-resident cells would be expected to result in equal rates of catabolism for all myelin lipids. We therefore determined whether individual lipids exhibit different turnover rates in purified myelin (Fig. 4C). Surprisingly, deuteration half-lives of lipids in purified myelin displayed a comparable level of heterogeneity to that seen in whole tissue, with very slow turnover of HexCer and very long chain SM (d42:2, d42:3) compared to PC and PI species, and intermediate turnover rates for PE O-/P- species (Fig. 4D). Differences in turnover rates were primarily a function of lipid class, however turnover rates for HexCer species decreased with the presence of a C-C double bond in the N-acyl chain, as observed in dissected corpus callosum. SHexCer species were turned over too slowly in isolated myelin for estimation of half-lives and 40/119 lipids showed significantly slower turnover in isolated myelin compared to corpus callosum tissue (Fig. 4E-J and Supplementary Data File 2).

### ApoE deficiency impairs the turnover of cholesterol and myelin sphingolipids

One mechanism through which different myelin lipids are replaced at different rates could be their differential binding to lipoproteins that transfer lipids to phagocytic cells. Since ApoE is one of the major lipid transport proteins in the brain, required for cholesterol clearance and remyelination following a demyelinating insult^35^, we tested whether ApoE deficiency perturbs physiological brain lipid turnover. ApoE deficiency significantly slowed the rate of decay in the weighted mean deuteration state for 36/114 lipids in the corpus callosum (Q<0.05) (Fig. 5Ai), of which cholesterol was most significantly affected (Fig. 5Bi), followed by HexCer (Fig. 5C), SHexCer (Fig. 5D), and PE O-/P- species (Supplementary Data File 3). The rate of cholesterol turnover was also slowed in the hippocampus of *Apoe-/-* mice (Fig. 5Aii,Bii). There was no effect of ApoE deficiency on turnover of individual lipid species in the cortex or isolated myelin, noting that statistical power was limited for the isolated myelin samples (n=3 per genotype) (Fig. 5Aiii-iv,Biii-iv). These findings indicate that ApoE regulates the turnover of cholesterol and other myelin-enriched lipids, a role that other lipoproteins cannot fully replace in its absence.

**Figure 5.**
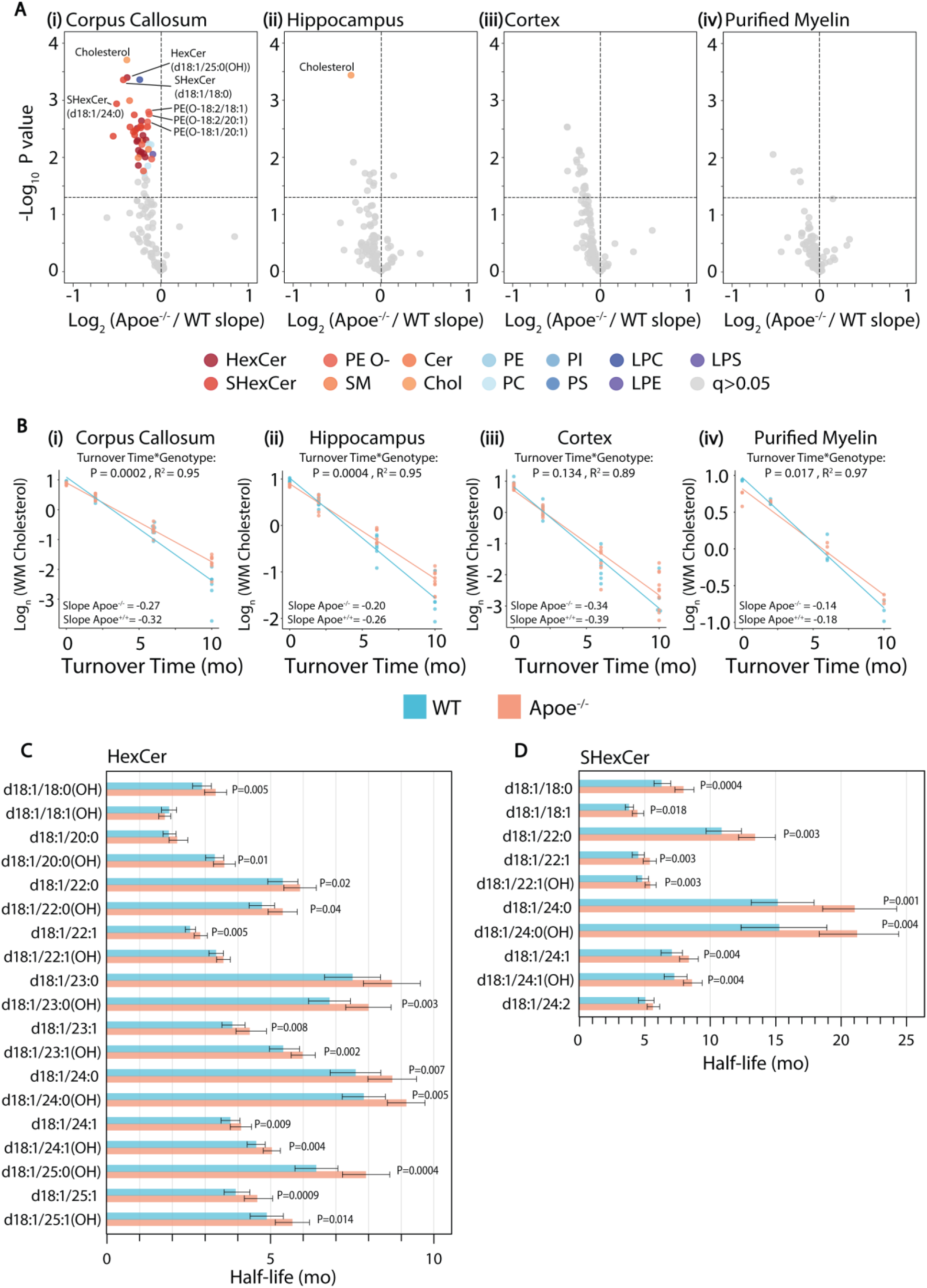
Impaired cholesterol turnover in *Apoe-/-* mice. (A) Volcano plots showing differences in lipid decay rates between *Apoe^-/-^* and WT mice in the corpus callosum (i), hippocampus (ii), cerebral cortex (iii), and purified myelin (iv). Exponential decay curves were linearised by natural log (ln) transformation and analysed by linear regression to assess turnover-genotype interactions. X-axes show log₂ fold-change in decay rates. (B) Ln-transformed decay curves for deuterated cholesterol in *Apoe^-/-^* and WT mice. Interaction P values and model fit (R²) are indicated. (C, D) Deuteration half-lives of individual HexCer (C) and SHexCer (D) species in the corpus callosum. Significant interactions between genotype and age (turnover time) are indicated with corresponding P values. The number and sex of mice in each group are provided in Extended Data Table 1 and Supplementary Data File 1.

### Myelin proteins have shorter half-lives than myelin sphingolipids and are not affected by ApoE deficiency

A key advantage with ^2^H_2_O administration is the unbiased labelling of both lipids and proteins in the same samples. Deuterium incorporation into tryptic peptides that map to myelin proteins, histones (known long-lived proteins)^27,28^ and cytoskeletal proteins was quantified, and used to determine their turnover rates (Fig. 6A-B). Deuteration half-lives were calculated as an average of three or more peptide half-lives, except for Cld11, for which the half-life of only one peptide could be calculated.

**Figure 6.**
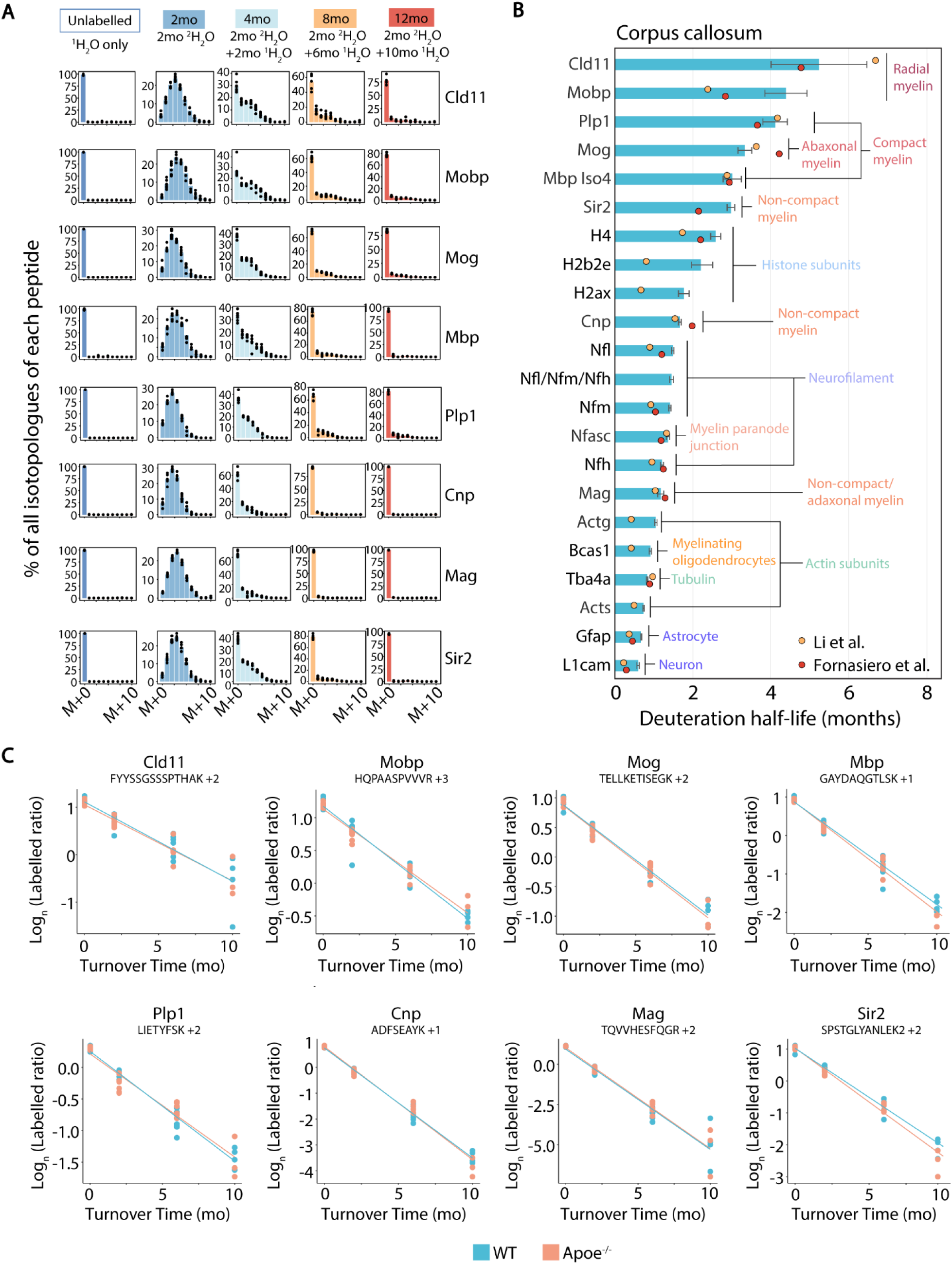
Myelin protein half-lives are unaffected by *Apoe* deficiency. (A) ^2^H isotopologue distributions for example peptides mapping to proteins of interest. (B) Mean protein half-lives (average half-life of multiple distinct peptides mapping to each protein) in WT mice. Error bars represent the 95% confidence interval of calculated half-lives. Cortical protein half-lives from published ^15^N labelling experiments are shown as orange^28^ and red dots^27^. (C) Decay curves (Ln-transformed labelled peptide ratio) for selected deuterated peptides that map to myelin proteins in *Apoe^-/-^* and WT mice.

Similar to studies that used ^15^N labelling to determine protein replacement rates^27,29^, radial and compact myelin proteins were turned over more slowly than histone subunits (H4, H2ax, H2b2e), neurofilaments, actin proteins, α4a tubulin, and Gfap (Fig. 6B). Proteins localised to radial (Cld11, Mobp) and compact (Plp1) myelin showed similar half-lives (∼4 months) to highly abundant HexCer species such as d18:1/24:1 (Fig. 2A). However, the half-lives for very long chain SHexCer and HexCer species, particularly those with saturated N-acyl chains, were much longer (7-15 months). Mog and Mbp (isoform 4), which are localised to the outer myelin layers and compact myelin, respectively, showed similar turnover rates to many PE species. In contrast, proteins localised to non-compact myelin (Cnp) or inner adaxonal layers of myelin (Mag) showed half-lives comparable to short-lived phospholipids (< 2 months). Overall, these results indicate that different proteins in the myelin sheath turn over at different rates, although the variation is less pronounced than seen with lipids. Unlike myelin-enriched lipids, the absence of ApoE did not have any effect on the half-lives of myelin proteins in the same mice (Fig. 6C), indicating that ApoE is required for turnover of long-lived myelin lipids, but not proteins, under normal physiological conditions.

### Myelin sphingolipid and cholesterol renewal decreases with ageing

To confirm that the differences in lipid decay rates seen with developmental ^2^H_2_O administration reflect rates of lipid replacement in adult mice rather than remodelling during development, we performed a complementary experiment in which wild-type (WT) or *Apoe-/-* mice were administered 25% ^2^H_2_O in their drinking water for 8 weeks at 3 or 12 months of age (Fig. 7A). Although the proportion of deuterated lipids was much lower, the number of ^2^H atoms per lipid was similar to that observed with the developmental labelling protocol (Extended Data Fig. 3), implying that the extent of deuteration reflects the rate of new lipid synthesis rather than substitution of ^1^H with ^2^H in existing lipids. New lipid incorporation was expressed as the percent deuteration for each lipid (Fig. 7B). In accordance with the rapid turnover rates of phospholipids in the developmental labelling protocol, all PI, PS, and most PC species in the corpus callosum showed over 80% replacement of non-deuterated with deuterated forms during the 8-week ^2^H_2_O administration period. PE and long chain SM species (d34:1, d36:0, d36:1) showed intermediate turnover rates, whereas the proportion of deuterated lipid molecules was 20-44% for HexCer, 6-22% for SHexCer and very long chain SM species, and 17% for cholesterol, indicative of slow replacement. The degree of HexCer and SHexCer deuteration was affected by N-acyl chain length and saturation, as observed with the developmental ^2^H_2_O administration protocol. Similarly, the proportion of deuterated lipid in isolated myelin was lowest for SHexCer, cholesterol, and very long chain SM species, and substantially lower for HexCer compared to glycerophospholipids (Extended Data Fig. 4). The deuterated proportion was lower for many lipids in isolated myelin compared to corpus callosum, further supporting our half-life estimates from developmental labelling.

**Figure 7.**
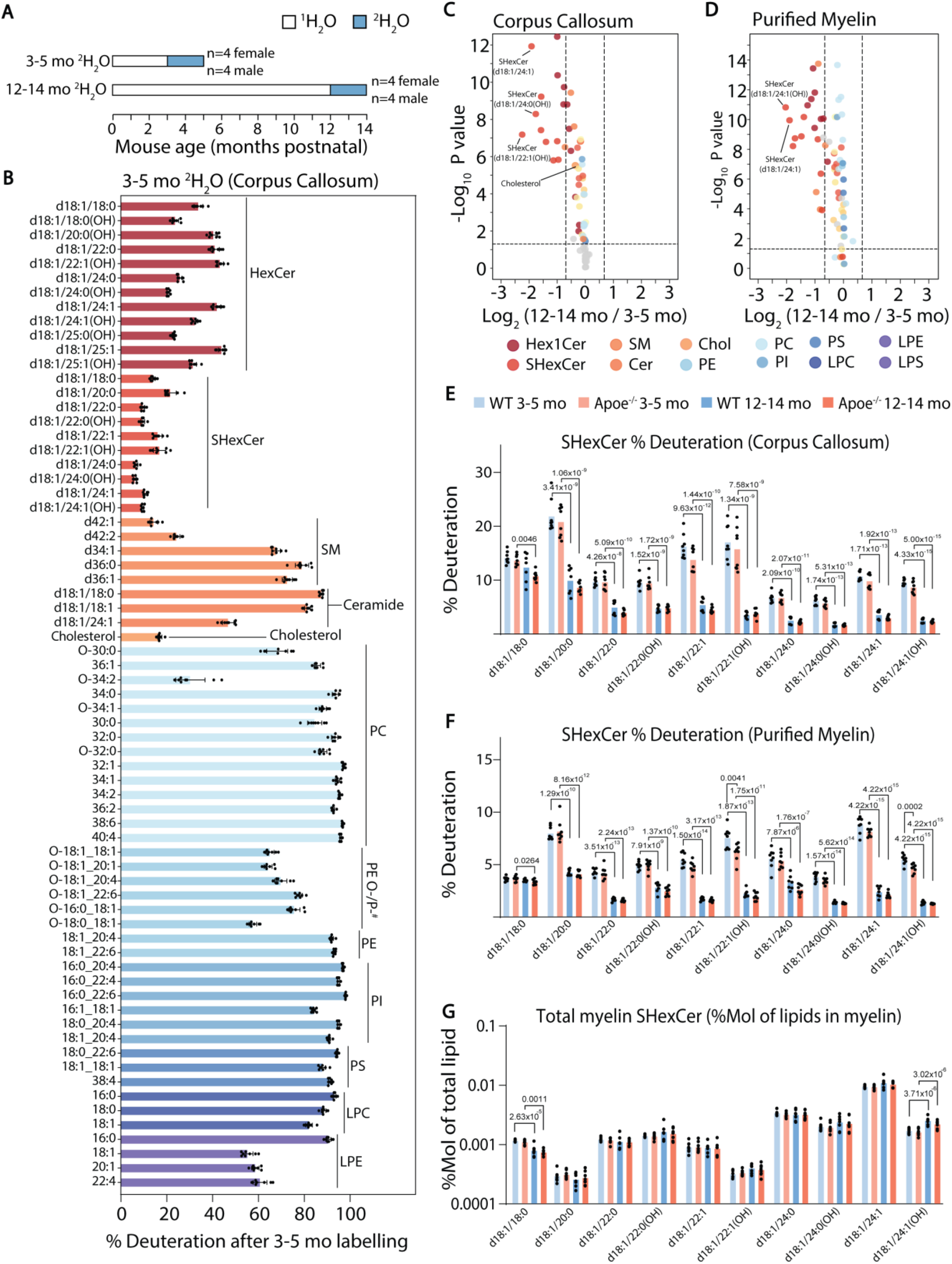
Myelin galactosphingolipid synthesis rates decline with ageing. (A) Schematic of ^2^H_2_O administration timeline in adult mice, with group sizes. (B) Deuterated proportion (%) for each lipid in the corpus callosum of WT mice following ^2^H_2_O administration from 3-5 months of age. (C-D) Volcano plots depicting differences in deuterated lipid proportion in (C) corpus callosum and (D) purified myelin of WT and *Apoe-/-* mice from the 12-14 month (n=15) compared to the 3-5 month (n=16) ^2^H_2_O administration group. Vertical dashed lines show a 1.5-fold difference, and horizontal line indicates P<0.05. Coloured datapoints are significant by two-way ANOVA after adjusting for multiple comparisons (Q<0.05). (E-F) Deuterated proportion for SHexCer species in the corpus callosum (E) and purified myelin (F) of WT and *Apoe-/-* mice administered ^2^H_2_O from 3-5 or 12-14 months of age. (G) Myelin SHexCer content as a percentage of total lipid in myelin. P values in E-G are derived from Tukey’s post-test, and are only shown for lipids that were significant at Q<0.05 for the genotype term in two-way ANOVA.

The effect of age and ApoE deficiency on lipid deuteration was assessed by two-way ANOVA (Supplementary Data Files 4 and 5). Deuteration of 52/69 lipids in corpus callosum and 60/64 in isolated myelin was significantly reduced in the 14-month old (12-month plus 8 weeks ^2^H_2_O) relative to the 5-month old (3-month plus 8 weeks ^2^H_2_O) mice (Fig. 7C,D), with the greatest reduction seen in SHexCer (Fig. 7E,F), followed by HexCer (Extended Data Fig. 5-6) species. Deuteration of several SM, PE P-/O-, and PC species was also modestly reduced in the corpus callosum of the 14-month compared to the 5-month-old mice, whereas PI species did not show any difference (Extended Data Fig. 5). In contrast, deuteration of almost all lipids in isolated myelin was decreased in the older mice, although the decrease was subtle for some (Extended Data Fig. 6). The reduction in deuterated proportion could be attributed to decreased addition of deuterated lipids to myelin over the 8 weeks of ^2^H_2_O administration (measured as the total amount of deuterated lipid in the myelin fraction) (Extended Data Fig. 7), whereas total lipid levels in myelin remained consistent for almost all lipids measured (Extended Data Fig. 8). Overall, these results demonstrate a reduction in the rate of myelin lipid turnover in the older mice, an effect that was pronounced for SHexCer. Despite this, the majority of SHexCer species did not differ as a proportion of the total lipid in myelin between the 14 and 5-months old groups (Fig. 7G), supporting a model in which their replacement rate in myelin is slower than that of other lipids, without affecting myelin lipid composition.

Cholesterol was most significantly affected by Apoe deficiency in both corpus callosum and isolated myelin of the adult-labelled mice (Extended Data Fig. 9 and Fig. 8A). Both age and ApoE deficiency caused a significant and additive reduction in the proportion of deuterated cholesterol in corpus callosum and purified myelin (Fig. 8B,C). Absolute levels of deuterated cholesterol did not differ in the corpus callosum of *Apoe-/-* compared to WT mice, indicating that the rate of cholesterol synthesis was not affected by ApoE deficiency (Fig. 8D). Instead, total cholesterol levels were 2-fold higher in the 14-month-old *Apoe-/-* mice (Fig. 8F), indicating that ApoE deficiency impairs cholesterol turnover. In contrast, levels of deuterated cholesterol were lower in isolated myelin from *Apoe-/-* compared to WT mice (Fig. 8E), while total cholesterol did not differ (Fig. 8G), indicating that ApoE deficiency retards the addition of newly-synthesised cholesterol to myelin.

**Figure 8.**
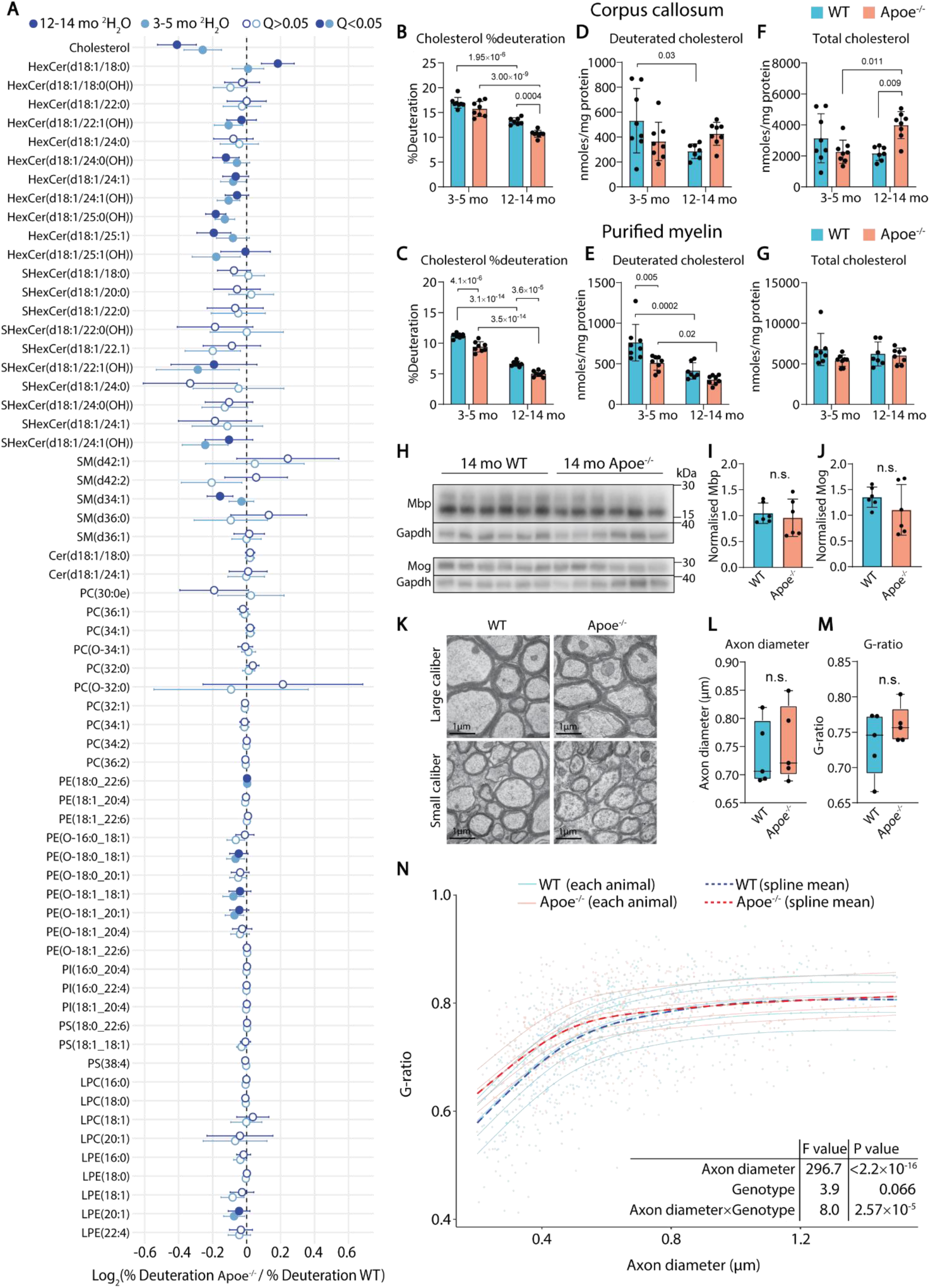
Addition of new cholesterol to myelin is ApoE-dependent and slows with ageing. (A) Forest plot depicting the change in % deuteration for all quantified lipids in purified myelin of *Apoe-/-* relative to WT mice. Mice were administered ^2^H_2_O from 3-5 or 12-14 months of age. Lipids significant for effect of genotype in 2-way ANOVA (Q<0.05) are shown as filled circles. (B,C) Proportion of cholesterol that is deuterated in the corpus callosum (B) and purified myelin (C). (D,E) Amount of deuterated cholesterol in the corpus callosum (D) and purified myelin (E). (F,G) Total cholesterol levels in the corpus callosum (F) and purified myelin (G). Error bars show 95% confidence intervals. (H) Western blots and (I, J) quantified intensity, relative to Gapdh, for Mbp (I) and Mog (J) in WT and *Apoe-/-* mice administered ^2^H_2_O from 12-14 months of age (6 mice per genotype). (K) Electron microscopy images of the corpus callosum of 12-month old WT and *Apoe-/-*mice. Fields containing smaller and larger calibre axons are shown for each genotype. (L, M) Mean corpus callosum axon diameter (L) and G-ratio (M) in each mouse (5 mice per genotype). (N) G-ratio as a function of axon diameter, including results of a linear mixed model used to determine the effects of axon diameter, genotype, and their interaction on G-ratio.

Western blotting for myelin proteins Mbp and Mog in the corpus callosum of 14-month-old *Apoe-/-*and WT mice showed no significant differences (Fig. 8H-J), suggesting that the effects of ApoE deficiency on lipid turnover and cholesterol levels are independent of gross myelin content. Similarly, measurements of myelin thickness (g-ratio) using electron microscopy (Fig. 8K) showed no significant differences between WT and *Apoe-/-* mice when considering axons of all diameters (Fig. 8L, M). However, myelin was significantly thinner (higher g-ratio) in small calibre axons (0.2-0.6 µm diameter) of *Apoe-/-* compared to WT mice (Fig. 8N), suggesting that the cholesterol trafficking defect in *Apoe-/-* mice impacts myelin maintenance.

## Discussion

This study describes a systems-level approach for quantification of lipid and protein turnover *in vivo*, establishing that brain lipid turnover rates are highly heterogeneous and primarily dependent on the headgroup. This heterogeneity in turnover rates was preserved in isolated myelin, where common PC and PI species exhibited half-lives of less than 2 months, whereas those of very long chain SM and SHexCer species exceeded 10 months. Myelin sphingolipid and cholesterol turnover slowed markedly from young adult (3 months) to post-reproductive (12 months) age, associated with reduced incorporation of newly-synthesised lipids into myelin; while disruption of extracellular lipid trafficking through ApoE deficiency preferentially impaired cholesterol turnover in the corpus callosum and the addition of newly-synthesised cholesterol to myelin. Our data indicates that distinct biochemical mechanisms regulate the turnover of different myelin components and provides direct evidence that myelin lipid renewal slows with ageing.

Myelin turnover under physiological conditions is thought to proceed through (i) the formation and degradation of myelinoid bodies^18^; (ii) phagocytosis of whole membrane sheaths^12,13^; and (iii) endocytosis by oligodendrocytes^18^. Blocking oligodendrocyte autophagy causes myelin protein accumulation and myelin thickening^12,13^, however lipid catabolism proceeds through endolysosomes rather than autophagosomes. Although our data indicates that different myelin constituents are turned over at different rates, this is not inconsistent with endocytosis as a mechanism of turnover. Our data supports a model in which glycerophospholipids, including PE plasmalogens, are more readily incorporated into degradative vesicles than sphingolipids and cholesterol, which localise to non-fluid myelin membrane raft domains containing Plp^36^. The parallel alignment of sphingolipid acyl chains, their interactions with cholesterol, and hydrogen bonding at the membrane surface contribute to the stability of these domains. Accordingly, we determined a half-life of ∼4 months for Plp, cholesterol, and abundant HexCer species in isolated myelin, whereas glycerophospholipids turned over more rapidly. This model aligns with the observation that sphingolipids with longer, saturated N-acyl chains had the longest half-lives, as the very long N-acyl chains of sphingolipids reduce membrane fluidity and are essential for myelin stability^37^, whereas C-C double bonds increase fluidity^37,38^.

It is possible that the extremely slow turnover of SHexCer reflects even less freedom of movement within the membrane, due to its localisation at myelin-axon adhesion sites. Although we determined a turnover rate for Nfasc overall, we were unable to calculate a turnover rate for the Nfasc155 isoform, since the detected peptides were common to multiple isoforms of Nfasc. SHexCer is thought to be required for stabilisation of myelin at the paranodes^39^ and our estimation of half-lives exceeding 4 months for SHexCer is consistent with the time-frame required to observe loss of sulfatides following inducible deletion of cerebroside sulfotransferase in adult mice^40^.

Another likely contributor to heterogeneous lipid turnover rates is their differential catabolism. Glycerophospholipids (PC, PE, PS, PI) can be catabolised by a wide range of cytosolic and extracellular phospholipases, whereas SHexCer and GalCer (HexCer) catabolism occurs in lysosomes, mediated by arylsulfatase A, galactocerebrosidase, and acid ceramidase^41^. It is possible that the susceptibility of polyunsaturated acyl chains in glycerophospholipids and the vinyl ether bond of plasmalogens to oxidation^42,43^ necessitates their more rapid turnover, compared to the more stable sphingolipids, in order to maintain myelin membrane integrity. Cytosolic, secreted, and inducible forms of phospholipase A_2_, which hydrolyses the *sn*-2 acyl chain of phospholipids, are expressed in oligodendrocytes following spinal cord injury^44^, and expression of the cytosolic and secreted forms following crush injury of the sciatic nerve is thought to facilitate myelin degradation^45^. The involvement of phospholipases in constitutive myelin phospholipid turnover remains largely unexplored and our ^2^H labelling approach provides a means to discover key mediators.

Cholesterol accounts for 40% of myelin lipid content by molarity^20^ and is essential for myelin formation and stabilisation^46,47^. The clearance and recycling of cholesterol in phagocytic cells is rate limiting for physiological myelin synthesis and remyelination following a demyelinating insult^35^. While the role of ApoE as a cholesterol transporter is well established^21,48^ and prior studies have shown that ApoE is required for cholesterol clearance from oligodendrocytes^49^ and phagocytic cells^35,50^, our study provides direct evidence that ApoE regulates the rate of cholesterol turnover in white matter and the rate of new cholesterol addition to myelin under normal physiological conditions. The reduced rate of new cholesterol incorporation into myelin in *Apoe-/-* mice may be a consequence of impaired cholesterol turnover, since the synthesis rate would need to be lowered to maintain correct myelin membrane composition. However, it is also possible that cholesterol incorporation into myelin membranes requires its transport by ApoE, possibly after synthesis in astrocytes^46^. The *APOE4* allele is the greatest genetic risk factor for dementia, and is associated with impaired cholesterol clearance from oligodendrocytes and reduced cortical myelin^49^. Future studies will investigate the effect of *APOE* genotype on cholesterol and myelin lipid turnover using *in vivo* ^2^H labelling.

ApoE deficiency also impaired the turnover of HexCer, SHexCer, and PE species, although with smaller effect size. The effect on SHexCer is consistent with prior work showing that SHexCer is carried on ApoE-containing CNS lipoproteins and accumulates in the brain of *Apoe-/-* mice^51^. These results indicate that ApoE is required for transport and degradation of myelin lipids, and this function cannot be entirely compensated by other lipoproteins. The requirement for ApoE in myelin lipid transport and importance of this process for myelin integrity very likely explain the significantly reduced myelin thickness on low calibre axons in ApoE deficient mice.

Ageing is associated with myelin thinning, segmental degeneration, reduced OPC differentiation, and impaired remyelination in both humans and mice^52^. Our observation that sphingolipid and cholesterol replacement slows significantly between 3 and 12 months of age suggests that renewal of compact myelin declines prior to overt structural change and is consistent with a prior study using a GFP reporter expressed only in newly-matured oligodendrocytes as a surrogate marker of new myelin synthesis^53^. A key advantage with ^2^H labelling is that it provides a quantitative read-out of endogenous lipid and protein turnover, revealing differences in turnover among different myelin constituents. Since our study only compared young adult to middle age in mice, it remains unclear whether this reduced rate of lipid renewal reflects a switch from plasticity to maintenance following myelin maturation throughout early adulthood, or a decline in myelin renewal that continues past middle age and contributes to the deterioration of white matter later in life, as suggested in other studies^53,54^.

Modern lipidomics has greatly advanced our understanding of the effect of genes and environment on physiology, and is a critical tool in diagnostic medicine^55,56^. However, measuring lipid levels at a snapshot in time is not sufficient to gain a mechanistic understanding of factors that control membrane homeostasis and the genetic basis of complex diseases^57^. The study demonstrates how combining stable isotope labelling with high resolution mass spectrometry will produce significant advances in our understanding of factors that regulate brain metabolism. A ^2^H_2_O administration study that predated the widespread use of LC-MS/MS for lipidomic analyses employed biochemical fractionation and gas chromatography-mass spectrometry to demonstrate that the total PC and PE pools in myelin are turned over faster than HexCer and cholesterol^33^. However, the authors could only determine aggregate turnover rates for major lipid classes, and estimates were based only on lipids that contained a palmitate (C16) acyl chain. In our study, high resolution LC-MS/MS allowed estimation of turnover rates for individual molecular species from a broader range of lipid classes, demonstrating that PE O-/P- species in myelin are turned over slower than PC or PI, but faster than HexCer and SHexCer; and that sphingolipid but not glycerophospholipid half-lives are affected by N-acyl chain length and the presence of C-C double bonds.

A limitation of our approach is that the deuteration half-lives reported herein may overestimate true lipid half-lives, due to recycling of deuterated fatty acids and lysolipids derived from lipid break-down. Although this will affect estimates of turnover rates in the developmental labelling paradigm, the relative lifetimes of different lipids were in strong agreement with the converse approach measuring deuteration of lipids in adult mice. Another advantage of the ^2^H_2_O labelling method is the ability to follow lipid and protein metabolism in the same samples. Our estimates of protein turnover rates using ^2^H_2_O administration were similar to those recorded with ^15^N labelling^27,28^, validating the utility of our approach for tracking protein turnover.

## Conclusion

Our findings indicate that myelin is turned over through a dynamic process whereby individual lipid and protein components are degraded and replaced at different rates, rather than removal and replacement of whole membrane sheaths. This constant turnover of myelin components is probably important for its long-term stability. The observation that myelin lipid renewal declines with ageing and is partially dependent on ApoE provides important mechanistic insight into why myelin integrity declines with ageing and how this process is accelerated by gene variants that are thought to confer dementia risk by altering brain lipid metabolism.

## Supporting information

Supplementary Data File 1 Deuteration decay half lives and LipidSearchID

Supplementary Data File 2 Linear modelling region differences in turnover

Supplementary Data File 3 Linear modelling APOE Age interaction

Supplementary Data File 4 Adult D2O CC ANOVA summary

Supplementary Data File 5 Adult D2O Myelin ANOVA summary

## Footnote

^#^Note that plasmalogen (plasmenyl) and plasmanyl ether lipid species cannot be distinguished using the mass spectrometry platforms used in this study, e.g. PE(P-34:1) and its mass isomer PE(O-34:2).

## Acknowledgements

We gratefully acknowledge subsidised access to the Sydney Mass Spectrometry, Laboratory Animal Services, and Sydney Microscopy and Microanalysis core facilities at the University of Sydney.

## Data Availability Statement

The raw lipidomic mass spectrometry data files will be made available through Zenodo upon acceptance. The proteomic dataset will be available on the PRIDE repository upon acceptance.

## Author Contribution statement

A.S.D. and J.Y.L. conceptualised and designed the study. J.Y.L. developed the analytical workflow, conducted developmental deuterium labelling experiments and performed data analysis for both labelling experiments. Y.C. conducted experiments and analysed data for the adult mouse labelling study, and performed western blotting. M.W., J.A.M., and S.R.E. acquired and analysed mass spectrometry imaging data. G.W., J.D.T., H.S., J.Y.L, and Y.C. performed mouse monitoring, euthanasia and brain dissections. J.D.T. acquired electron microscopy data. J.D.T., G.W. and J.Y.L. analysed electron microscopy data. A.S.D. and S.R.E. acquired funding. J.Y.L. and A.S.D. wrote the manuscript, with input from all authors.

---

**Extended Data Fig. 1.**
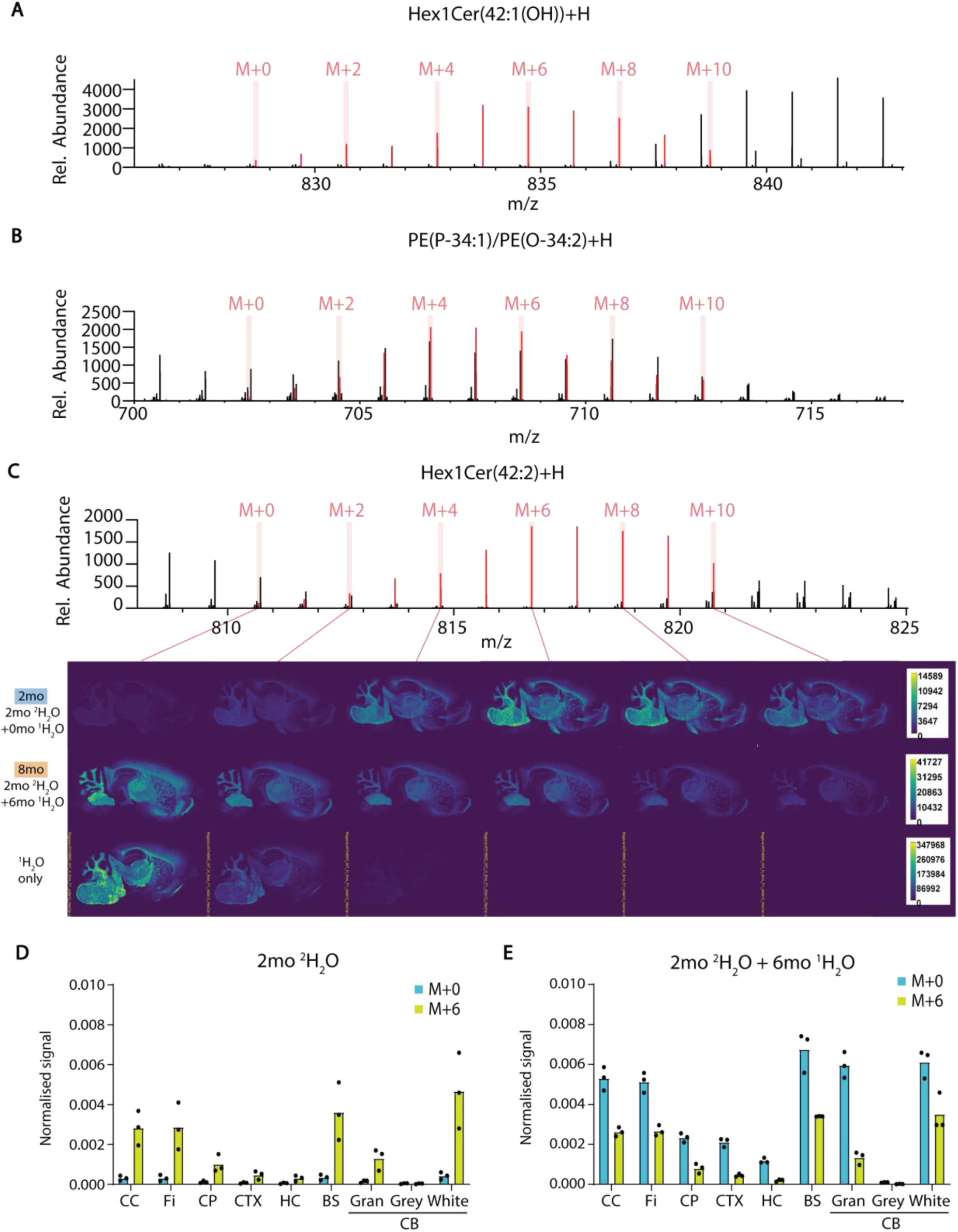
(A, B) Mass spectra for (A) [HexCer(d42:1(OH))+H]^+^ and (B) [PE(P-34:1)/PE(O-34:2)+H]^+^, averaged over the section. Mass-to-charge (*m/z*) values corresponding to the non-deuterated (M+0) and deuterated (M+2 – M+10) isotopologues of each lipid ion are shown in red. (C) Representative mass spectrum and corresponding ion images for [HexCer(d42:2)+H]^+^. (D, E) Quantified M+0 and M+6 ion intensities for [HexCer(d42:2)+H]^+^, normalised to the total ion chromatogram at each pixel. Normalised signals were averaged over defined regions: CC, Corpus callosum; Fi, fimbria; CP, caudate putamen; CTX, cerebral cortex; HC, hippocampus; BS, brain stem; CB, cerebellum; Gran, cerebellar granular layer; Grey, cerebellar grey matter; White, cerebellar white matter. Bars show the mean (3 mice per group).

**Extended Data Fig. 2.**
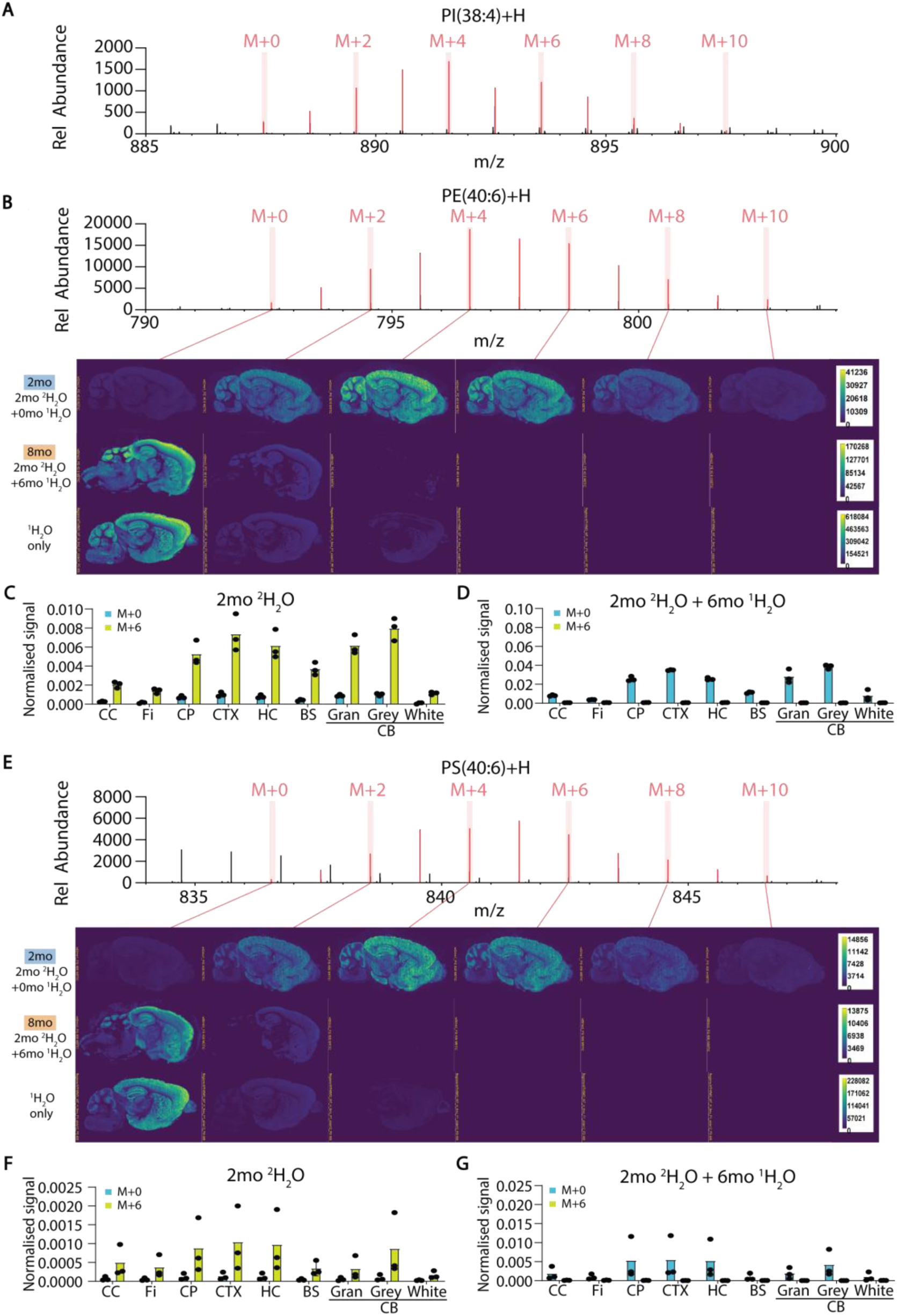
(A) Raw mass spectra for [PI(38:4)+H]⁺. (B) Raw mass spectra and corresponding ion images for [PE(40:6)+H]⁺. (C-D) Quantification of M+0 and M+6 isotopologues of [PE(40:6)+H]⁺, normalised to the total ion chromatogram (TIC) at each pixel. (E) Raw mass spectra and corresponding ion images for [PS(40:6)+H]⁺. (F-G) Quantification of M+0 and M+6 isotopologues of [PS(40:6)+H]⁺, normalised to the TIC at each pixel. Normalised signals were averaged over defined brain regions: CC, corpus callosum; Fi, fimbria; CP, caudate putamen; CTX, cerebral cortex; HC, hippocampus; BS, brain stem; CB, cerebellum; Gran, cerebellar granular layer; Grey, cerebellar grey matter; White, cerebellar white matter. Bars show the mean (3 mice per group).

**Extended Data Fig. 3.**
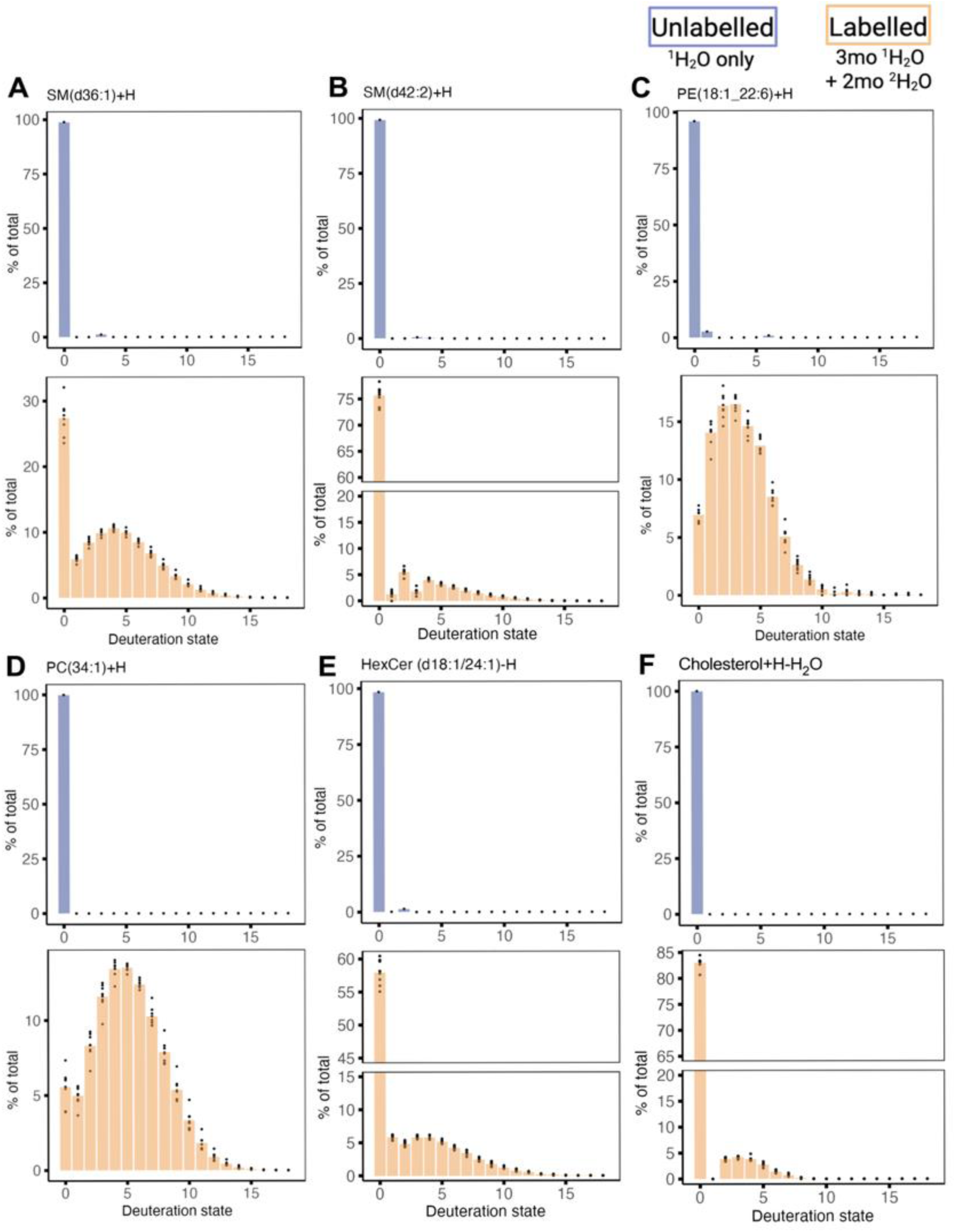
Isotopologue distribution patterns for (a) SM(d36:1), (b) SM(d42:2), (c) PE(18:1_22:6), (d) PC(34:1), (e) HexCer(d18:1/24:1), and (f) cholesterol in the corpus callosum of mice administered ^2^H_2_O from 3-5 months of age (orange), or mice given only normal drinking water (blue).

**Extended Data Fig. 4.**
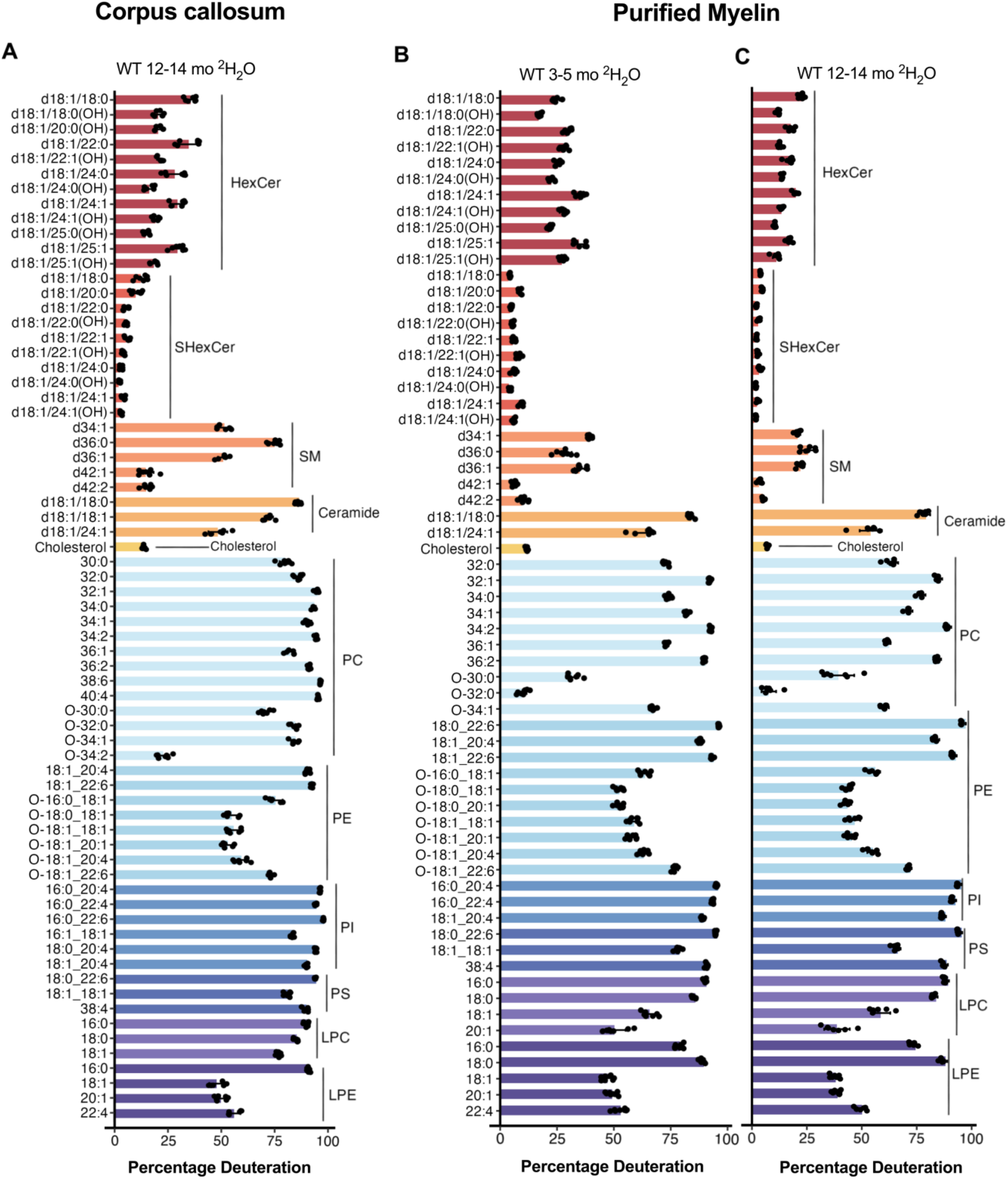
Percentage deuteration of lipids in (A) corpus callosum of WT mice administered ^2^H_2_O from 3-5 months of age (n=7), (B) purified myelin of WT mice administered ^2^H_2_O from 3-5 months of age (n=8), and (C) purified myelin of WT mice administered ^2^H_2_O from 12-14 months of age (n=7).

**Extended Data Fig. 5.**
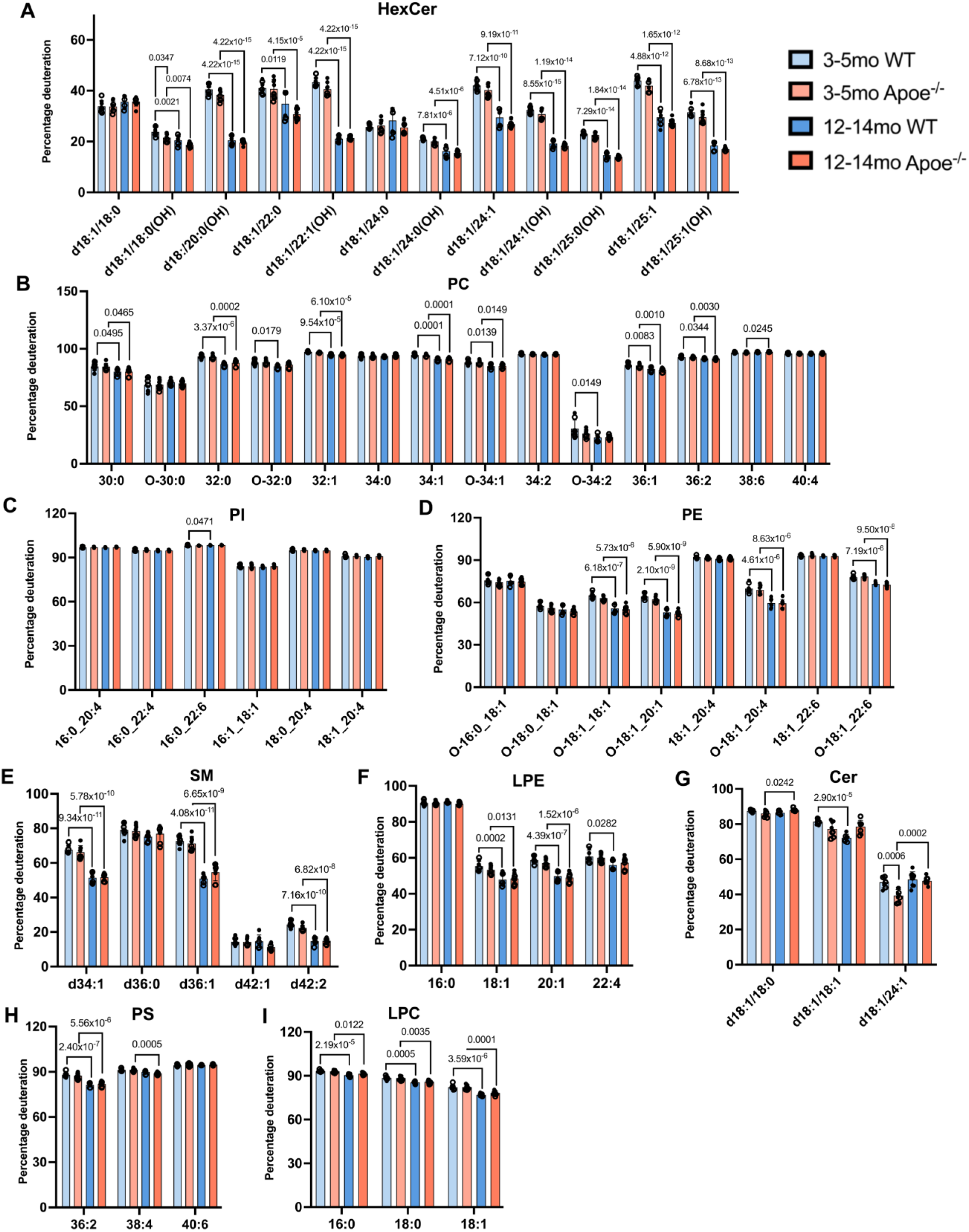
Percentage deuteration of (A) HexCer, (B) PC, (C) PI, (D) PE, (E) SM, (F) LPE, (G) Ceramide, (H) PS and (I) LPC in corpus callosum of WT and *Apoe^-/-^* mice administered ^2^H_2_O from 3-5 or 12-14 months of age. P values for Tukey’s post-test comparisons are shown for lipids significant in 2-way ANOVA.

**Extended Data Fig. 6.**
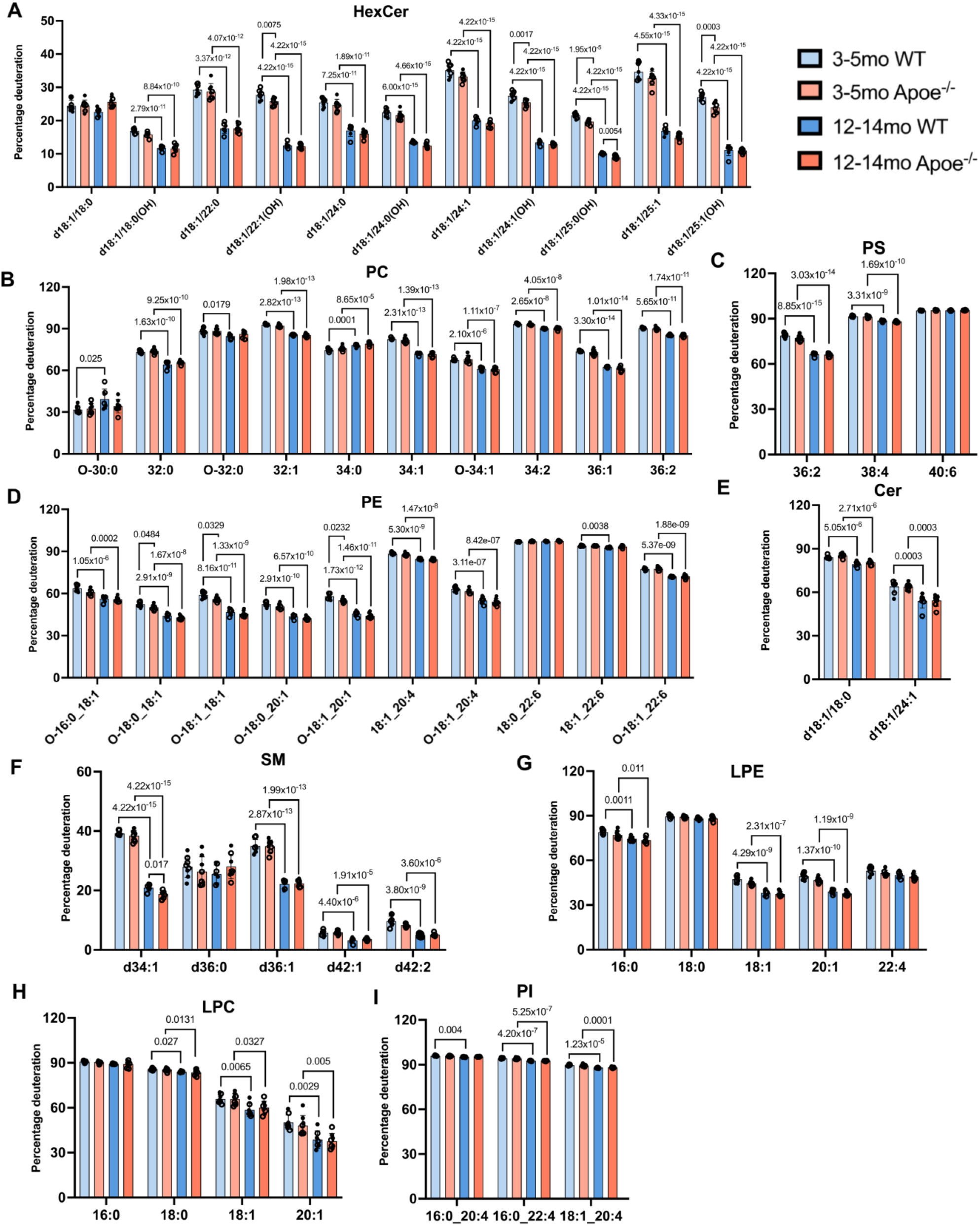
Percentage deuteration of (A) HexCer, (B) PC, (C) PS, (D) PE, (E) Ceramide, (F) SM, (G) LPE, (H) LPC, and (I) PI in purified myelin of WT or Apoe^-/-^ mice administered ^2^H_2_O from 3-5 or 12-14 months of age. P values for Tukey’s post-test comparisons are shown for lipids significant in 2-way ANOVA.

**Extended Data Fig. 7.**
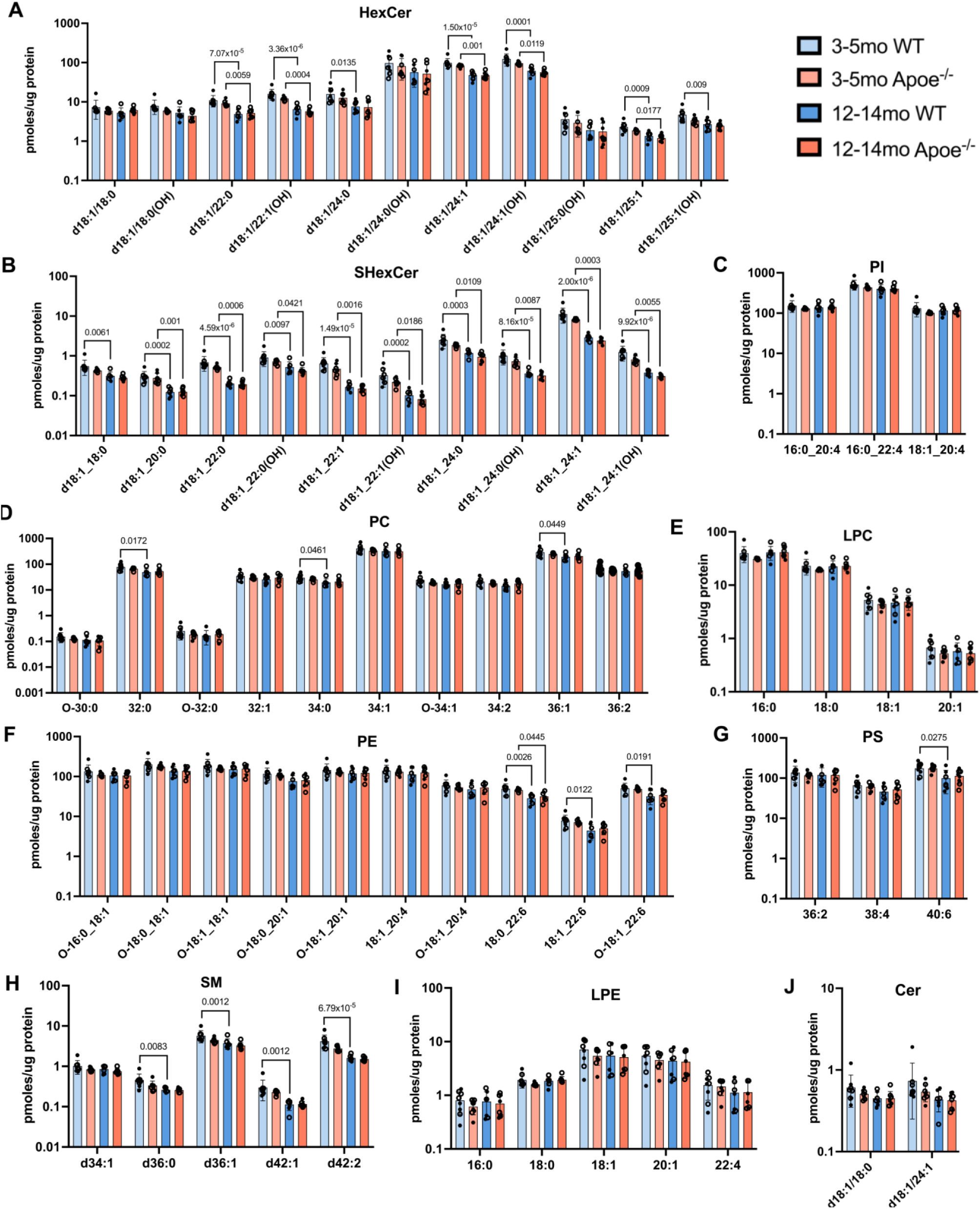
Total level of deuterated (A) HexCer, (B) SHexCer, (C) PI, (D) PC, (E) LPC, (F) PE, (G) PS, (H) SM, (I) LPE and (J) Ceramide in purified myelin of WT and Apoe^-/-^ mice administered ^2^H_2_O from 3-5 or 12-14 months of age. P values for Tukey’s post-test comparisons are shown for lipids significant in 2-way ANOVA.

**Extended Data Fig. 8.**
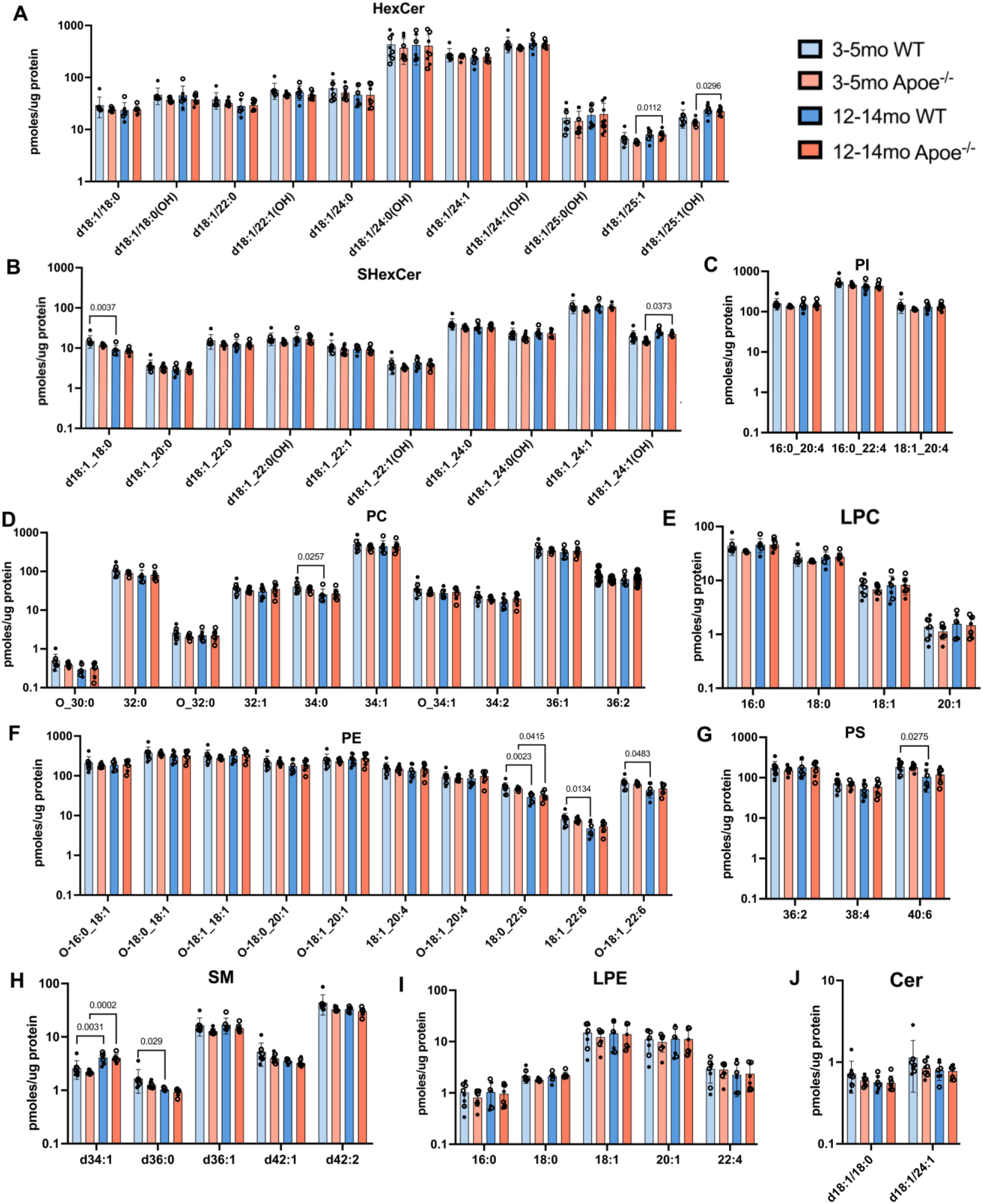
Total level of (A) HexCer, (B) SHexCer, (C) PI, (D) PC, (E) LPC, (F) PE, (G) PS, (H) SM, (I) LPE and (J) Ceramide in purified myelin of WT and Apoe^-/-^ mice administered ^2^H_2_O from 3-5 or 12-14 months of age. P values for Tukey’s post-test comparisons are shown for lipids significant in 2-way ANOVA.

**Extended Data Fig. 9.**
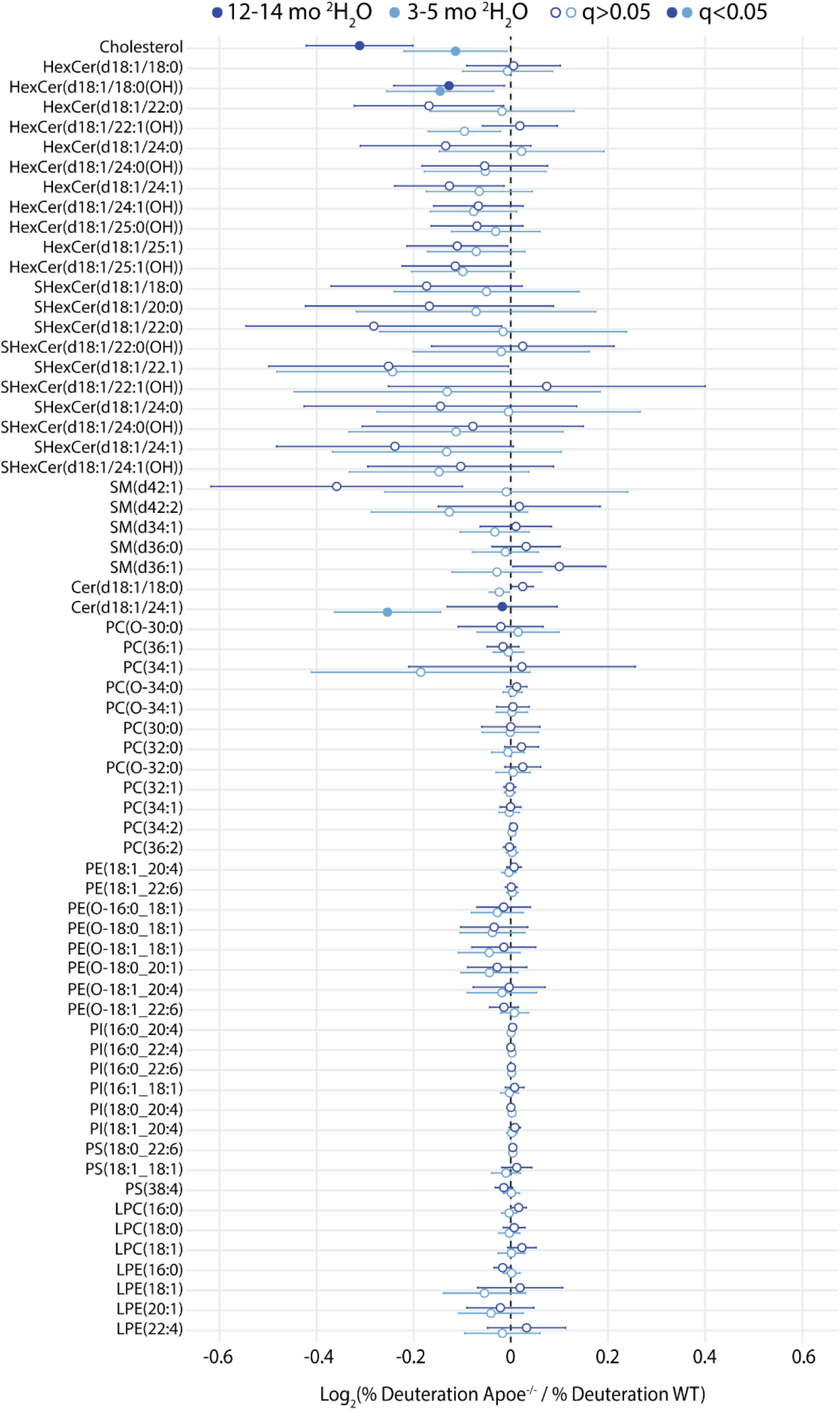
Forest plot showing the change in % deuteration for lipids in the corpus callosum of *Apoe-/-* relative to WT mice. Mice were administered ^2^H_2_O from 3-5 or 12-14 months of age. Lipids significant for effect of genotype in 2-way ANOVA (q<0.05) are shown as filled circles.

## Methods

### Deuterium oxide administration to mice

Experiments were conducted in accordance with the Australian code of practice for the care and use of animals for scientific purposes and approved by the University of Sydney animal ethics committee (#2022-2136). Mice were housed in groups of four with a 12 h light/dark cycle, in plastic cages with straw nesting, a cardboard roll and wooden chew sticks, and given food and water ad libitum.

#### Developmental deuterium oxide (^2^H_2_O) administration

^2^H_2_O was purchased from Merck (cat# 151882-4KG) and filter-sterilised prior to mixing with drinking water. Time-mated pregnant female wild-type (WT) C57BL/6J and *Apoe*-/- mice (C57BL/6J background) were shipped from Animal Resource Centre (Perth, Australia) at day 12 of pregnancy. Pregnant WT and *Apoe-/-* dams, 3 months of age, were given 15% ^2^H_2_O in their drinking water from gestational day 14 (E14) until they gave birth. The mother and her pups were kept on 15% ^2^H_2_O until post-natal day 7 (P7), when the ^2^H_2_O was increased to 20%. At P13, the ^2^H_2_O concentration was increased to 25%, and remained at this concentration until 2 months of age (P60), at which point all mice were switched to normal drinking water (^1^H_2_O). Mice were culled at 2 months (i.e. 2 months ^2^H_2_O only), 4 months (2 months ^2^H_2_O + 2 months ^1^H_2_O), 8 months (2 months ^2^H_2_O + 6 months ^1^H_2_O), and 12 months (2 months ^2^H_2_O + 10 months ^1^H_2_O) of age. Mice were monitored and weighed twice weekly, with no signs of distress or toxicity observed. All mice were euthanised with isofluorane, then transcardially perfused with saline prior to removal of the brain for dissection. Corpus callosum, hippocampus, and cerebral cortex were dissected from one hemisphere and stored at -80 °C. The other hemisphere was either snap frozen for myelin isolation or slow frozen over dry ice and stored at -80 °C for mass spectrometry imaging. Mouse numbers used for corpus callosum (CC) and hippocampus samples are shown in Supplementary Table 1. An additional five *Apoe-/-* mice (one 2-month, two 8-month, and two 12-month females) were available at the time of analysis of cortex samples. One sample each for CC, cortex, and hippocampus was detected as an outlier in analysis of the LC-MS/MS data and excluded; and one hippocampus sample (WT, 12 month) was excluded due to inaccurate dissection. Two- and 8-month-old C57BL/6J mice given only normal drinking water were used as controls for lipid identification.

**Supplementary Table 1.** Mouse numbers for developmental ^2^H_2_O administration protocol.

| Age (months) | WT male | WT female | *Apoe*<sup>-/-</sup> male | *Apoe*<sup>-/-</sup> female |
| --- | --- | --- | --- | --- |
| 2 | 3 | 3 | 4 | 2 |
| 4 | 3 | 5 | 4 | 3 |
| 8 | 2 | 5 | 3 | 3 |
| 12 | 3 | 4 | 8 | 2 |

#### Adult mouse ^2^H_2_O administration

Two-month-old WT C57BL/6J and *Apoe*-/- mice were purchased from Animal Resource Centre (Perth, Australia). Groups of 4 male and 4 female mice were administered 25% ^2^H_2_O in their drinking water for 8 weeks at 3 or 12 months of age. One 12-month-old male WT mouse was euthanised prematurely and excluded from data analysis due to excessive weight loss and abnormal circling behaviour. Mice were euthanised with isoflurane on the final day of the 8 week ^2^H_2_O administration, perfused with saline, brains removed, and one hemisphere was dissected as described above. The other hemisphere was snap frozen for myelin isolation.

### Preparation of brain tissue homogenates

Crude tissue homogenates were prepared by ultrasonicating the dissected tissue in 500 µL of 20 mM Hepes pH 7.4, 10 mM KCl, EDTA-free cOmplete protease inhibitor cocktail (Roche), 1 mM dithiothreitol, and 3 mM β-glycerophosphate, for 5 min (30 s on/30 s off) at 4 °C in a Qsonica Q800R2 sonicating bath. Total protein concentrations were determined using the Bradford assay (Bio-Rad).

### Myelin isolation

Myelin sheaths were enriched from frozen half brains using sucrose density centrifugation^58^. Brain tissue was homogenised in 0.3 M sucrose solution (20 mM Tris-Cl pH 7.45, 2 mM Na_2_EDTA, 1 mM DTT, cOmplete protease inhibitor, Roche) using 20-30 Eppendorf micropestle strokes. Homogenates were then layered over 0.83 M sucrose and ultracentrifuged for 30 min at 75,000×g, 4 °C. Crude myelin that formed at the interface was collected, resuspended in Tris-buffered solution (20 mM Tris-Cl pH 7.45, 2 mM Na_2_EDTA, 1 mM DTT, cOmplete protease inhibitor, Roche), and washed twice by centrifugation; once at 75,000×g and once at 12,000×g. The resulting crude myelin pellets were then resuspended in 0.3 M sucrose and taken through another round of gradient centrifugation over 0.83 M sucrose as described above. Further purification of resulting myelin pellets was performed by resuspending in 0.83 M sucrose, prior to layering 0.3 M sucrose over the myelin for ultracentrifugation. The resulting pure myelin interface was collected, resuspended in Tris buffer, and spun down to pellet purified myelin. Myelin pellets were resuspended in Tris buffer and stored at - 80 °C for LC-MS/MS analysis and western blotting.

### Lipid extraction

For the developmental labelling paradigm, lipids were extracted from mouse tissue homogenates and purified myelin samples (∼100 µg protein and ∼20 µg protein, respectively) using two rounds of single phase butanol-methanol (BUME) (1:1 v/v) extraction^59^. In brief, sample homogenates were mixed with 1.2 mL of BUME and sonicated for 1 h at 20 °C (30s on/off), with 80% amplitude, in a Qsonica Q800R2 sonicating bath. The samples were then centrifuged at 2000×g, 20 °C, to pellet any insoluble precipitate, and the lipid-containing liquid phase was transferred to a glass tube. The pellet was resuspended in an additional 1 mL of BUME, vortexed, and sonicated for 1 h. Following another centrifugation step using the same settings as previously, the supernatant was added to the original glass tube. The solvents were evaporated overnight at RT in a Thermo Fisher SpeedVac SC210, after which the lipids were reconstituted in 80% methanol:water and transferred to glass HPLC vials with 300 µL fused inserts for immediate analysis. Samples were stored at 4 °C prior to analysis.

For the adult labelling paradigm, lipids were extracted using two rounds of two-phase extraction with methyl-tert-butyl ether (MTBE) and methanol. Sample homogenates were mixed with 250 µL of methanol and 850 µL of MTBE, and sonicated in a Qsonica Q800R2 sonicating bath for 30 min at 80% amplitude at 4 °C (30s on/off). Phase separation was induced by adding 100 µL of MilliQ water (adjusted based on homogenate volume), followed by centrifugation at 2000×g for 5 min, at 20 °C. The upper organic phase containing lipids was transferred into a glass tube, and the remaining aqueous phase was re-extracted with the addition of 150 µL methanol, 500 µL MTBE and 125 µL MiliQ water. The samples were sonicated, vortexed and centrifuged under the same conditions as the first round of extraction, and the upper phase was collected and pooled with the initial extract. The solvents were evaporated in a Thermo Fisher SpeedVac SC210 (overnight at RT), after which lipids were reconstituted in 50% butanol-methanol (1:1 v/v) and transferred to glass HPLC vials with fused inserts.

For the adult mouse labelling experiments, the following internal standards were added at the start of lipid extraction: 30 nmoles of β-sitosterol; 5 nmoles of 19:0/19:0 phosphatidylcholine; 4 nmoles of 14:0/14:0/14:0/14:0 cardiolipin; 2.5 nmoles d18:1/17:0 sulfatide; 2 nmoles each of d18:1/17:0 sphingomyelin, d18:1/17:0 glucosylceramide, 17:0/17:0 phosphatidylserine, 17:0/17:0 phosphatidylethanolamine, 17:0/17:0/17:0 triglyceride, 17:0/17:0 phosphatidylglycerol and 17:0 cholesteryl ester; 1 nmole each of 17:0/17:0 phosphatidic acid, d18:1/15:0-d7 phosphatidylinositol, d18:1/15:0-d7 diacylglycerol; 0.5 nmoles each of d18:1/17:0 ceramide, 17:1 lysophosphatidylethanolamine, 17:1 lysophosphatidylserine, 17:0 lysophosphatidylcholine, 18:1-d7 monoacylglycerol, and d18:1/12:0 dihexosylceramide; and 0.2 nmoles each of 17:1 sphingosine, 17:1 sphingosine 1-phosphate, d3-16:0 acylcarnitine, and 17:0 lysophosphatidic acid.

### Lipidomic analysis using liquid chromatography-tandem mass spectrometry (LC-MS/MS)

Lipids were detected on a Thermo Fisher Q-Exactive HF-X mass spectrometer with Vanquish HPLC (ThermoFisher Scientific). A Waters Acquity UPLC CSH 2.1 × 100 mm C18 column (1.7 μm particle size) was used, with flow rate 0.28 ml/min. Mobile phases were A: 10 mM ammonium formate, 0.1% formic acid, 60% acetonitrile and 40% water; B: 10 mM ammonium formate, 0.1% formic acid, 10% acetonitrile and 90% isopropanol. Run time was 25 min using a binary gradient starting at 20% B for 3 min, increasing to 45% B from 3 to 5.5 min, then to 65% B from 5.5 to 8 min, then to 85% B from 8 to 13 min, then to 100% B from 13 to 14 min. The gradient was held at 100% B from 14 to 20 min, then decreased to 20% B and held to 25 min. Precursor ion data was acquired in full scan/data-dependent MS^2^ mode (resolution 120,000 FWHM at 200 *m/z*, scan range 220-1600 *m/z*). The ten most abundant ions in each cycle were subjected to MS^2^, with an isolation window of 1.4 *m/z*, collision energy 30 eV, resolution 17,500 FWHM, maximum integration time 110 ms and dynamic exclusion window 10 s. An exclusion list of background ions was based on a solvent blank.

### Deuterated lipid isotopologue detection and distribution analysis

Lipid extracts from mice administered ^2^H_2_O were run concurrently with those from mice given only ^1^H_2_O (unlabelled mice). Lipids in unlabelled samples were identified through matched precursor (mass tolerance 5 ppm) and product (mass tolerance 8 ppm) ions using LipidSearch software (version 4.2, ThermoFisher) (Supplementary Data File 6). The list of identified lipids was then used to create a list of theoretical precursor *m/z* and retention time values for each isotopologue from M+0 (no deuterium atoms) to M+18 (18 deuterium atoms), based on an increase in *m/z* of 1.006276746 for each deuterium (^2^H). Peak areas for this list were quantified using Skyline software (version 23.1), with mass tolerance 10 ppm and retention time tolerance +/- 0.3 min, after which the theoretical abundance of naturally occurring isotopes was subtracted using the IsoCor package (ver 2.2.2)^60^ for Python (ver 3.11.8). Peak areas for each deuterated isotopologue were then expressed as a percentage of the total abundance of all isotopologues for a given lipid. Lipids showing co-eluting peaks at *m/z* corresponding to a deuterated isotopologue in samples from non-deuterated (^1^H_2_O only) control mice after isotope correction were excluded from further analysis, applying the rule that the M+0 peak must be >95% of the total peak area for each lipid in the non-deuterated control samples. In the developmental labelling experiment, the distribution of isotopologues was expressed as the weighted mean deuteration state, according to the formula: 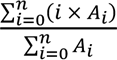, where *i* indicates the n^th^ isotopologue number (the number of deuterium atoms incorporated) and *A_i_* the peak area of that isotopologue.

In the adult labelling experiment, deuteration was expressed as a percentage of the total peak area for that lipid, comprising the deuterated + non-deuterated forms after isotope correction, i.e. deuterated/(deuterated + non-deuterated). Absolute amounts of non-deuterated and deuterated lipid in each sample were estimated as the (peak area for each lipid isotopologue/peak area for the class-specific internal standard) × amount of internal standard added, then normalised to protein concentration as estimated by Bicinchoninic acid assay (Thermo Fisher Scientific #23225).

### Western blotting

Samples containing 5 µg protein in lithium dodecyl sulfate (LDS) sample buffer (ThermoFisher, # NP0007) plus reducing reagent (ThermoFisher, #NP0009) were heated at 85 °C for 10 min, resolved at 150 V for 50 min on 4-12% Bis-Tris Plus polyacrylamide gels (ThermoFisher, #NW04120BOX), and transferred onto polyvinylidene difluoride (PVDF) membrane (ThermoFisher, #88518) at 20 V for 1 h. Membranes were blocked with 5% skim milk in 50 mM Tris, pH 7.6, 150 mM NaCl, and 0.1% Tween-20 (TBST) for 1 h at RT, and washed 3 × 5 min with TBST. Membranes were probed with primary antibody in TBST overnight at 4 °C and washed with TBST before incubation with anti-rabbit IgG HRP or anti-mouse IgG-HRP secondary antibody at RT for 1 h. Bands were detected using enhanced chemiluminescence (ECL) reagent (ThermoFisher Scientific, #32109), imaged using ChemiDoc Touch Imaging System, and quantified with BioRad ImageLab 6.1. Primary antibodies used were: rabbit anti-Calnexin (Abcam #ab22595, RRID #AB_2069006), mouse anti-CNPase (Abcam #ab6319, RRID #AB_2082593), mouse anti-COX IV (Cell Signalling #11967, RRID #AB_2797784), rabbit anti-GAPDH (Cell Signalling #2118, RRID #AB_561053), rabbit anti-Olig 2 (Abcam #ab109186, RRID #AB_10861310), rabbit anti-Synaptophysin (Abcam #ab32127, RRID #AB_2286949), rabbit anti-Myelin Basic Protein (Abcam #ab40390, RRID #AB_1141521), rabbit anti-PLP (Abcam #ab28486, RRID # AB_776593), mouse anti-GFAP (Cell Signalling #3670, RRID #AB_561049), rabbit anti-Myelin oligodendrocyte glycoprotein (Abcam #ab32760, RRID #AB_2145529), rabbit anti-Iba1 (Abcam #ab178847, RRID #AB_2832244), mouse anti-βIII tubulin (Bio-Legend #801202, RRID AB_2313773). All antibodies were used at 1:3000 dilution with the exception of anti-βIII tubulin, which was used at 1:5000. Novex Sharp pre-stained protein ladder was used to mark molecular weights (ThermoFisher, #LC5800).

### Label-free proteomics

Crude homogenate (10 µg protein) was extracted with 100 µL 4% SDS, 100 mM NaCl, 20 mM NaPO_4_ (pH 6), 10 mM NaF, 10 mM tris(2-carboxyethyl)phosphine (TCEP), 10 mM N-ethylmaleimide (NEM), 10 mM sodium pyrophosphate, 2 mM sodium orthovanadate, 60 mM sodium β-glycerophosphate, and EDTA-free cOmplete protease inhibitor cocktail (Roche). The volume was made up to 150 µL using MilliQ water and samples were incubated for 10 min at 65 °C with shaking (500 rpm), then ultrasonicated for 10 min at 20 °C (15s on/15s off). Proteins were precipitated with chloroform/methanol/water in the ratio 1:4:1:3 (sample:methanol:chloroform:water)^61^, dried down, and reconstituted in 30 µL of 8 M Urea in 0.1 M Tris-HCl (pH 8.0). Protein concentrations were measured by BCA assay (ThermoFisher Scientific), after which protein samples were diluted 8-fold in 0.1 M Tris-HCl (pH 8)/1 mM CaCl_2_ and digested for 16 h at 37 °C with 0.1 µg trypsin (#90058, ThermoFisher). Digestion was stopped with the addition of trifluoroacetic acid to a final concentration of 1%, and the samples were centrifuged to pellet any undigested protein (18,000×g, 10 min, 20 °C). The supernatants were transferred to new tubes and subjected to solid-phase extraction^62^. Data was acquired using data-independent acquisition on a ThermoFisher Scientific Q-Exactive HF-X mass spectrometer coupled to an EASY-nLC system, using previously described chromatography and mass spectrometer parameters^63^. Samples were randomised in their injection order and unlabelled samples were injected every 5 deuterium-labelled samples.

### Deuterated peptide isotopologue distribution analysis

Proteins were identified in unlabelled (^1^H_2_O only) mouse samples by a library-free search using DIA-NN (version 1.9.1) with the Uniprot mouse database downloaded on 27/08/2024. The spectral library generated from the search was then used to perform a peptide search in both labelled and unlabelled samples (using DIA-NN), based on the expectation that the naturally occurring isotopologues from unlabelled samples will elute at similar retention times to the M+n peptide isotopologues of labelled samples. Importantly, samples with high relative M+0 isotopologue abundance for peptides allow for accurate peptide identification and retention time inference in extensively deuterated samples.

Output .speclib files were then imported into Skyline (version 23.1) to visualise peak quality in unlabelled mice (^1^H_2_O only). A list of *m/z* and matched retention times for peptides mapping to proteins of interest was used to generate theoretical isotopologue *m/z* for each peptide (M+0 to M+15) using an in-house R script. Theoretical isotopologue masses were reimported into Skyline for the quantification and visualisation of isotopologues, constrained to the same retention time (±0.5 min). Peptide isotopologues were corrected for the presence of naturally occurring isotopes using IsoCor (ver 2.2.2) and expressed as a percentage of the total abundance of all isotopologues for a given peptide. Peptides were excluded from analysis if isotopologue peaks (greater than 5% abundance) were observed in samples from unlabelled mice following isotope correction.

Deuteration was expressed as a fraction of the total peak area for each peptide, comprising the deuterated + non-deuterated forms after isotope correction. The deuterated fraction was used to determine peptide turnover rates, as per previous studies using ^13^C-lysine or ^15^N instead of ^2^H^27,28^. Peptide half-lives determined from the deuterated fraction were very similar to those calculated with the weighted mean deuteration state, except that the weighted mean produced higher variance. Protein half-lives were estimated as an average of the half-lives of three or more distinct peptides from that protein.

### Mass spectrometry imaging (MSI)

Frozen brain hemispheres were sectioned along the sagittal plane (15 µm thickness) at -20°C using a CM1950 cryo-microtome (Leica Microsystems Pty Ltd, Victoria, Australia), mounted onto glass slides and stored at -80°C until analysis. Sections at sagittal levels 10-13 were used for MSI. Following removal from -80°C tissue sections were brought to room temperature for 15 min in a vacuum desiccator, then dipped in refrigerated 100 mM aqueous ammonium fluoride solution (Sigma-Aldrich, Castle Hill, NSW, Australia) for 3 x 5 s, dried again for 60 min in a vacuum desiccator, and coated using 20 mg of 2,5-dihydroxyacetophenone (Sigma-Aldrich, Castle Hill, NSW, Australia) matrix using an in-house sublimation system at 140 °C for 2.5 min. Following sublimation, samples were recrystalised in a chamber for 90 s at 50 °C with 1 mL of 0.5% ethanol and stored in a vacuum desiccator until imaging.

MSI was performed using a prototype timsTOF Pro mass spectrometer (Bruker Daltonics, Bremen, Germany). The system includes an atmospheric pressure matrix-assisted laser desorption ionisation (MALDI) ion source combined with a plasma post-ionisation system (SICRIT, Plasmion GmbH, Augsburg, Germany) and has been described in detail previously^64^. MSI was performed in positive ion mode at a pixel size of 30 × 30 µm^2^ using beam scanning dimensions of 15 × 15 µm^2^. The laser was manually focussed onto the sample and operated at a repetition rate of 5 kHz with 200 laser shots accumulated at each position. Desorbed species were collected in the heated inlet capillary (middle temperature of 360°C) and transferred to the SICRIT device for plasma post-ionisation which was operated at 1600 V amplitude and 15000 Hz frequency.

### Hematoxylin and Eosin Staining

Sections were stained with hematoxylin and eosin after MSI. Matrix was removed with 3 x 30 s washes in 100% methanol, and sections were rehydrated stepwise with 2 x 2 min 95% ethanol, 2 x 2 min 75% ethanol, and 2 min deionised water washes. Sections were stained with Harris hematoxylin for 3 min, washed with running tap water until clear, then washed for 1 min in deionised water. Eosin was applied for 30 s and samples were dehydrated with 95% ethanol for 2 x 30 s, then 100% ethanol for 2 x 30 s, then xylene for 30 s before coverslipping with DPX mount and #1.5 coverslip glass. Images were acquired using a Leica SP8 FALCON (Leica Microsystems CMS GmbH, Mannheim, Germany) with 10x objective.

### MSI Data Analysis

All data was processed using SCiLS Lab 2025 (13.01.17218, Bruker Daltonics, Bremen, Germany). Hematoxylin/eosin images were manually annotated in QuPath^65^ v0.5.1 to define regions of interest, which were imported into SCiLS Lab using the annotation plugin for QuPath. Given the moderate resolution of the orthogonal TOF mass analyser and the complex spectra generated, many isotopologues likely have unresolved isobaric interferences, e.g. M+2 deuterated lipid ions cannot be readily distinguished from naturally-occurring isotopes of lipid ions with one less double bond (i.e. type II isobaric interference). Therefore, the lipids discussed in this paper were carefully selected based on inspection for minimal isobaric interference and spatial correlation between isotopologues. Average peak areas within a selected *m/z* ± 14 ppm from the annotated regions were exported as a CSV file and visualised as stem plots using MATLAB (R2024b, MathWorks). Typical mass errors were less than 5 ppm. The ion images (un-normalised) for selected *m/z* values were extracted with the SCiLS application programming interface (2025b) using Python 3.13, normalised to total ion current on a per-pixel basis, then converted using NumPy (2.0.2) into CSV files for visualisation using Fiji image analysis software (Image J)^66^. Python was used to construct boxplots of the data using the Seaborn (0.13.2) data visualisation library and NumPy 2.0.2.

### Electron Microscopy

Mice were anaesthetised with isofluorane and trans-cardially perfused with 0.9% heparinised saline, followed by 2.5% glutaraldehyde/1% paraformaldehyde in phosphate buffered saline (PBS). The brains were removed and post-fixed overnight at 4°C in the same fixative. The solution was replaced with 30% sucrose in PBS and the tissue was stored at 4°C until use. Fixed brain tissue was sectioned along the sagittal plane at roughly 2 mm thickness. Using a brain punch kit, biopsy punches were acquired along the corpus callosum. The biopsies were washed in PBS and underwent secondary fixation in 1% osmium tetroxide in PBS, followed by dehydration through 30% to 100% ethanol solutions. The samples were infiltrated and embedded in medium epon resin. Ultra-thin 70 nm sections were acquired on a Leica Ultracut 7 using a diamond knife and collected on copper grids. Sections were contrast stained using 2% uranyl acetate and Reynold’s lead citrate, and images were captured on a transmission electron microscope (FEI Tecnai T12). Using ImageJ, the axonal lumen diameter and the outer diameter of myelin surrounding the axons were measured perpendicular to the longest axis and the g-ratio was calculated by dividing the axon lumen diameter by the outer diameter of the axon.

Differences in the mean axon diameter and myelin thickness were determined by Welch’s t-test. To determine whether the relationship between axon diameter and myelin thickness differed between genotypes, a repeated-measures linear mixed-effects model was used to account for multiple axons measured within each mouse and avoid pseudo-replication. Axons in the range 0.2 to 1.5 µm were used for this analysis to focus on biologically relevant diameters and avoid effects from sparsely sampled extremes. Axon diameter, genotype, and their interaction were included as variables. Significance for each term was determined using Type III ANOVA with Satterthwaite’s approximation (as applied in R lmer/lmerTest).

### Statistical analyses

All data and statistical analyses were performed using R Studio. Unless otherwise indicated, P values were adjusted for multiple comparisons using the Benjamini-Hochberg false discovery rate (FDR) correction at 5% (i.e. q < 0.05 was considered significant). The normal distribution of residuals for all statistical analyses were assessed using the Anderson-Darling normality test and the Shapiro-Wilk normality test. In cases where the test indicated non-normally distributed residuals, QQ plots and histograms were used to assess the degree of deviation from normality to decide whether variables should be natural log-transformed to fulfil the assumptions of normal distribution. For regression analyses of deuterated lipid turnover, which follow exponential decay kinetics, strict normality of residuals is not expected even after log-transformation due to bounded measurements and heteroscedasticity across timepoints. In these cases, the appropriateness of the model was evaluated based on linearity after log-transformation and homoscedasticity of residuals.

The change in weighted mean deuteration state (ΔWM) relative to the 2-month timepoint was 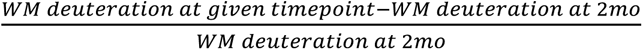 and expressed as as percentage. These values were fit to exponential decay curves for all 220 lipid ions, of which 182 ions showed good fit (R^2^>0.9). Exponential decay curves were used to determine deuteration half-lives and 95% confidence intervals for each lipid ion. In cases where multiple ions were detected for the same lipid, e.g. -H and +HCOO ions of HexCer species, deuteration half-life values were averaged.

Differences in lipid class turnover rates between brain regions were assessed using repeated-measures ANOVA, treating each lipid as an independent datapoint within its class. Differences in turnover rates of individual lipids between brain regions or between WT (Apoe⁺^/^⁺) and Apoe⁻^/^⁻ mice were assessed by linear regression. For these analyses, weighted mean intensities of a representative ion for each lipid were natural log-transformed to linearise exponential decay, and models tested the interaction between age and region or genotype, with significant interaction terms indicating differences in turnover rate (slope). Lipids for which weighted mean values approached zero in more than 50% of samples per group (by region or genotype at each timepoint) were excluded to avoid invalid log transformations.

The effect of age (3 or 12 months) and genotype (WT or *Apoe^-/-^*) on % lipid deuteration and total levels of each lipid was assessed by two-way ANOVA with Tukey’s post-test, correcting main effect p-values for FDR (Supplementary Data File 4 and 5). Graphs show p-values resulting from the post-tests, but only for lipids significant at q<0.05 for main effect of age, genotype, or age-genotype interaction.

